# Pathways in the brain, heart and lung influenced by SARS-CoV-2 NSP6 and SARS-CoV-2 regulated miRNAs: an *in silico* study hinting cancer incidence

**DOI:** 10.1101/2024.02.11.578752

**Authors:** Shrabonti Chatterjee, Joydeep Mahata, Suneel Kateriya, Gireesh Anirudhan

## Abstract

The influence of SARS-CoV-2 non-structural protein in the host’s tissue-specific complexities remains a mystery and needs more in-depth attention because of COVID-19 recurrence and long COVID. Here we investigated the influence of SARS-CoV-2 transmembrane protein NSP6 (Non-structural protein 6) in three major organs – the brain, heart, and lung *in silico*. To elucidate the interplay between NSP6 and host proteins, we analyzed the protein-protein interaction network of proteins regulated after SARS-CoV-2 infection and that are interacting with NSP6 interacting proteins. Pathway enrichment analyses provided global insights into biological pathways governed by differentially regulated genes in the three tissues after COVID-19 infection. Many drugs targeting hub genes of tissue-specific protein interactome were found that could be candidates for COVID-19 management. MiRNA-gene network for the tissue-specific regulated proteins was also deduced and comparing gene list targeted by SARS-CoV-2 regulated miRNAs, we found three and two common genes in the brain and lung respectively. Among the five common proteins revealed as potential therapeutic targets across the three tissues, Galectin3 (LGALS3) that was upregulated in the heart and brain after COVID-19 infection is reported to be influencing all the ten hallmarks of cancer positively and is found in multiple cancers. COVID-19 infection also causes myocardial inflammation and heart failure (HF). HF is observed to be increasing cancer incidence. Our bioinformatics and systems study hints probable effect of COVID-19 infection in cancer incidence and warrants in-depth studies in this direction and cancer surveillance especially with the present scenario of long COVID-19 and recurrent COVID-19 infections.

**Graphical Abstract:** 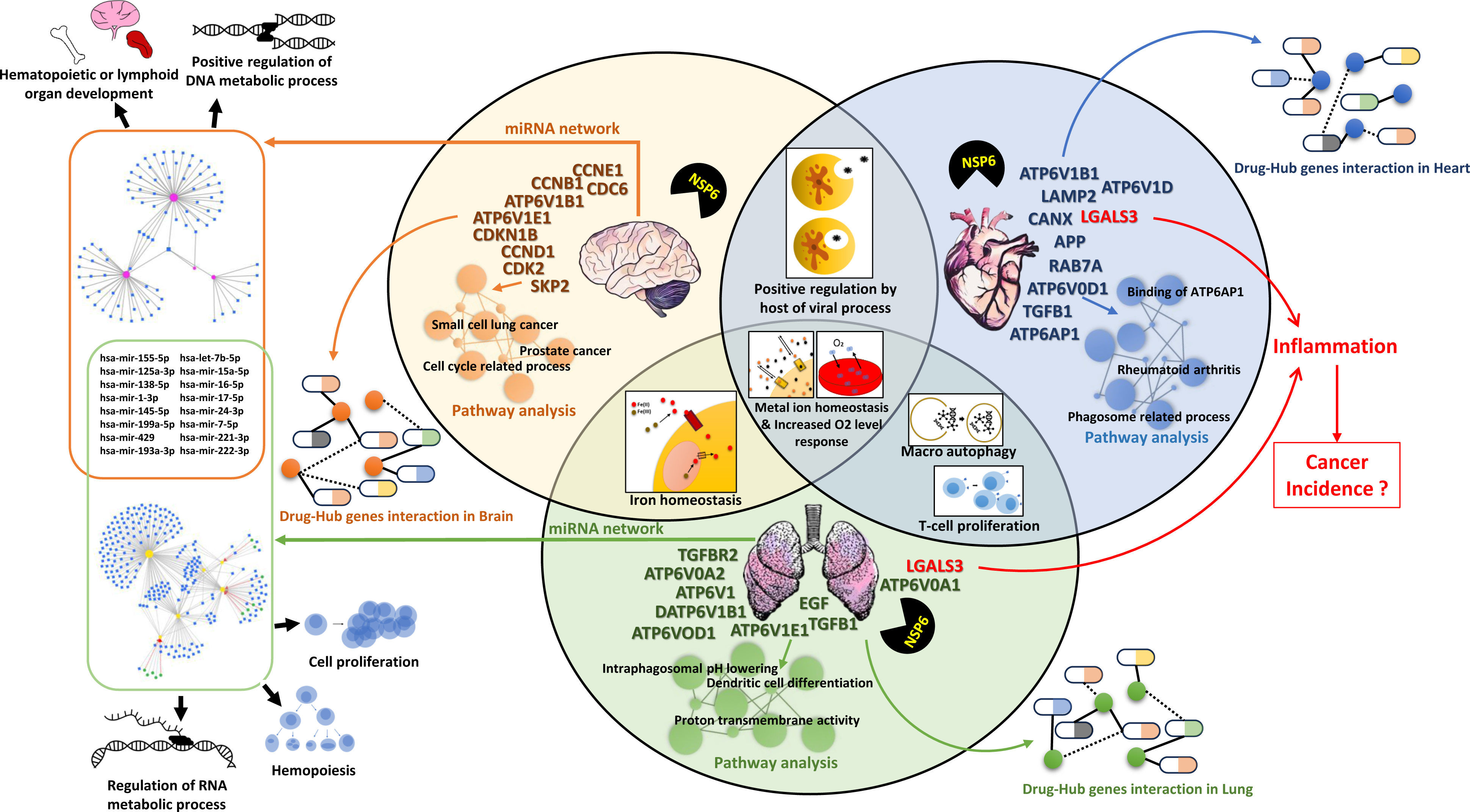

## 1. Introduction

After the disastrous COVID-19 pandemic, we are forced to live along with COVID-19 as a new normal. The unavailability of specific drugs that directly target COVID-19, the ineffectiveness of vaccines against emerging variants and long COVID are some of the prominent responsible factors. In addition, reports of various animals as hosts,^1–8^ emergence of variants and detection of these clades in humans coming in contact with them,^9,10^ also increase the risks of emergence of zoonotic variants. Changes in the symptoms of COVID-19 and decreased routine testing add to the present troubles in the identification of infected individuals and there by COVID-19 spread. Multiple organ systems get impacted due to COVID-19^11^ in the acute phase of the disease and several symptoms can persist from many weeks to two years,^12^ now termed as “Post-acute sequelae of COVID-19” (PASC).^13^ Increased risk of PASC including stroke, and various cardiac disorders, after post COVID-19 as well as it’s reinfection are well documented.^12–21^ In addition, other risks include chronic kidney diseases, fatigue, muscle weakness, acute coronary disease, memory problems, acute kidney injury, thromboembolism, constipation, gastro-oesophageal reflux disease (GERD), etc.^12^ In post COVID-19 cases, increased incidence of stroke^22^ and cardiovascular diseases^23^ even in patients who did not require hospitalization prompts us to study the probable mechanistic pathways affecting these organs in comparison to the prominent primary target organ, the lungs.

Various COVID-19 proteins are shown to be interacting with different host proteins.^24,25^ Multiple proteins coded in the SARS-CoV-2 genome can be mainly divided into structural proteins, nonstructural proteins (NSPs) and accessory proteins.^26–28^ 16 NSPs are encoded in ORF1a and ORF1b that have multiple enzymatic functions.^28^ Functions include regulation of viral RNA replication, transcription, and genome replication.^28^ One of the NSP proteins NSP6, formed by the cleavage of ORF1a polyprotein by NSP5 with chymotrypsin-like cysteine protease (3CLpro) activity contains 290 aa.^28,29^ Predicted structure using Alphafold proposed “one C-terminal, two anti-parallel beta-sheets, sixteen turns and fourteen α helices”.^30^ Regarding the transmembrane domains, different reports suggest transmembrane helices as six,^29,31^ seven,^32^ and eight.^30^ NSP6 is present in both SARS-CoV and SARS-CoV-2. Amino acid sequence alignment among these shows sequence identity of 88.2% and sequence similarity of 98.3%.^33^ SARS-CoV-2 NSP6 was found to be homodimerizing by both fluorescence resonance energy transfer (FRET) and co-immunoprecipitation studies.^29^ The C-terminal domain with part of amphipathic helix and dimerisation were found to be involved in endoplasmic reticulum (ER) remodelling.^23^ NSP6 is reported to be involved in SARS-CoV-2 replication by restricting ER luminal proteins accessing double membrane vesicles (DMVs) but allowing lipid flow, positioning and organising DMV clusters, and facilitating lipid droplets (LDs) through LD-tethering complex DFCP1-RAB18.^29^ Observation of emerging deletion mutation of NSP6 (Δ106–108) in different variants of SARS-CoV-2 like Alpha, Beta, Gamma, Eta, Iota and Lambda^29,34^ and its gain of function with a higher ER-zippering activity as reported by Ricciardi et al.,^29^ increases its significance to be studied further. Omicron’s attenuated virulence was also shown to be influenced by mutations in S and NSP6.^35^ Antagonism of type-I interferon (IFN-I) signaling by the enhanced suppression of STAT1 and STAT2 phosphorylation by mutant SARS-CoV-2 NSP6 with triple deletion (ΔSGF or ΔLSG) was observed.^36^ Two proteins ATP6AP1 and Sigma–1–receptor were reported to be interacting with NSP6 and are also drug targets.^24^ Sigma–1–receptor ligands were observed to be inhibiting SARS-CoV-2 infection in Vero E6 cell line.^24^ Overexpression of NSP6, NSP8, and M in stem cell differentiated cardiomyocytes showed altered transcriptome and proteome, and induced apoptosis.^37^ Effect of NSP6 in SARS-CoV-2 replication through properly formed NSP6 connectors and lipid droplets also underscores its significance.^29^ These observed functional significance of NSP6 prompts us to study various significant pathways influenced by it and putative drug targets for the viral containment.

In the present *in silico* work, we analyzed to look for pathways influenced by NSP6 in three tissues, the brain, heart and lung. Among proteins interacting with SARS-CoV-2 NSP6 interacting proteins expressed in these tissues, those influenced by SARS-CoV-2 infection (18, 45 and 23 interacting proteins in the brain, heart and lung respectively) were analyzed to find out pathways and biological processes involved by these proteins. In the brain, heart and lung there were 25, 147 and 168 pathways respectively. To find out the prominent players, hub genes were found and drugs (approved and non-approved) targeting these were searched for in the Drug-Gene Interaction Database (DGIdb). Next, we wanted to find out the influence of miRNAs in these proteins and hence putative miRNA players were searched that could influence the proteins of our interest. In addition, reported SARS-CoV-2 influenced miRNAs were probed for their targets in the interacting proteins of SARS-CoV-2 NSP6 interacting protein list influenced by SARS-CoV-2 infection in the brain and lung. Reported SARS-CoV-2 regulated circulating miRNAs targeting SARS-CoV-2 regulated proteins in the heart, interacting with SARS-CoV-2 NSP6 interacting proteins were used to find out prominent miRNA players in the heart during SARS-CoV-2 infection. All deduced pathways and prominent players could be helpful in finding out druggble pathways/targets for better management of SARS-CoV-2 infection.

## 2. Materials and methods

### 2.1. SARS-CoV-2 NSP6 interacting proteins

To find out SARS-CoV-2 NSP6 influenced pathways in the brain, heart and lungs, we took a list of proteins interacting with SARS-CoV-2 NSP6.^24^ These four reported interacting proteins (SIGMAR1, ATP13A3, ATP5MG, ATP6AP1) were considered for further analyses.

### 2.2. Tissue-specific genes interacting with NSP6 interacting proteins

For finding the interacting proteins of SARS-CoV-2 NSP6 interacting proteins in a tissue-specific manner, IID (Integrated Interactions Database) (http://ophid.utoronto.ca/iid) was used based on two or more experimental pieces of evidence (studies or bioassays).^38^ Common genes present in the gene list obtained from IID and genes regulated after SARS-CoV-2 infection in the brain choroid plexus epithelium,^39^ heart,^40^ and lung^41^ were used for further analyses. Data obtained were from the Log_2_FC values from these published data. Brain choroid plexus epithelium was infected with SARS-CoV-2 isolate that was sequenced^42^ from the first patient reported in USA for COVID-19 who was cured.^43^ Heart specimens were from an autopsy cohort consisting patients who underwent intermittent mandatory ventilation (IMV) periods (0-30 days) from symptom appearance to death.^40^ Lung samples were from two deceased COVID-19 human subjects and 2 normal lung samples were obtained post-surgery.^41,44^ We considered genes having the p-value of ≤ 0.05 for our further analyses. To find out the common and unique genes from different tissues, we used jvenn.^45^

### 2.3. Construction of tissue-specific protein-protein-interaction network of SARS-CoV-2 NSP6 influenced proteins

Tissue-specific and SARS-CoV-2 NSP6 influenced proteins were used to make protein-protein-interaction network using IID. Common genes present in the gene list obtained from IID and genes regulated after SARS-CoV-2 infection in the brain choroid plexus epithelium,^39^ heart,^46^ and lung^41^ were used. In these common genes, those with p-value ≤ 0.05 were selected. Further we discerned up– and downregulated genes, that had a cut off value of log_2_ fold change ± 2.0. These categorized gene lists were poised for subsequent in-depth investigation and analyses.

### 2.4. Identification of central hub genes in tissue-specific protein-protein interactome

The tissue-specific genes were the input for the construction of protein–protein interaction network in STRING version 12.0^47^ with default settings (Network Type: full STRING network, Required score: medium confidence (0.400), FDR stringency: medium (5 percent)). The network was further grown to obtain more tissue-specific interactions and these were uploaded in Cytoscape version 3.10.1.^48^ Hub genes for the tissue-specific network were found using cytoHubba plugin.^49^ To initiate the hub gene identification process, a Node Score was calculated, employing the Maximum Clique Centrality (MCC) metric. The MCC metric, the most accurate among the topological analysis methods,^49^ focuses on identifying cliques which is a subset of nodes within a network. Based on the node’s centrality the MCC calculates the size of the maximum clique by providing rank for a specific node. Subsequently, an analysis was executed, employing the criteria of the top 10 hub genes within the designated PPI network. Further refinement was carried out by selecting parameters pertaining to the first stage node, shortest path, and expanded network analysis. These steps collectively led to the delineation of the hub gene network.

### 2.5. Gene ontology and KEGG pathway analysis of central hub genes in tissue-specific protein-protein interactome

Subsequent to ranking the top 10 hub genes using the MCC method, we proceeded with a comprehensive pathway analysis. For this purpose, we employed ClueGO, an additional plugin available in the Cytoscape software suite.^50^ Using the functional analysis mode, we conducted targeted searches, with default medium network specificity settings and a pathway significance threshold set at ≤ 0.05 for p-values. With the help of ClueGO, GO terms of the identified hub genes network were based on Gene Ontology (GO) Biological Process, GO Cellular Component, GO Molecular Function, GO Immune System Processes, Reactome pathways, and Kyoto Encyclopedia of Genes and Genomes (KEGG) pathways. Kappa score threshold was taken as 0.4 and for statistical analysis, we used Enrichment/Depletion (two-sided hyper geometric test). Kappa score also known as Cohen’s coefficient, provides inter-relatability between two functional groups based on the genes shared by them.

### 2.6. Drug-gene interaction analysis of hub genes in DGIdb

To find out possibility of detected hub genes as probable drug targets, we searched for drugs targeting these genes using the Drug-Gene Interaction database and further identified the gene categories for the probable prognostic hub genes (DGIdb v.5.0.1, https://www.dgidb.org/).^51,52^ Interactions of drug-gene network and gene categories were uploaded in Cytoscape version 3.10.1.^48^ to construct network.

### 2.7. Tissue-specific pathway enrichment analysis

To obtain the Gene Ontology (GO) IDs for particular tissues, we utilized the R package ClusterProfiler.^53^ We executed this process individually for the genes associated with the brain, heart, and lung tissues. This enabled us to extract pathways specific to each tissue. For a more comprehensive analysis, we further categorized the genes into upregulated and downregulated groups. We performed separate analyses on the genes exhibiting upregulation and downregulation to find the biological processes and pathways associated with them. Subsequently, we employed a significance threshold by setting a q-value cutoff of 0.05 and only these significant pathways were considered for dot-plot construction by using ggplot2 package in R.^54^

### 2.8. Tissue-specific miRNA Network Construction

NSP6 may regulate the activities of a diverse array of miRNAs. miRNAs are well-known influencers of gene expression. Using miRNet (https://www.mirnet.ca),^55^ we have identified and constructed a tissue-specific interaction network of SARS-CoV-2 influenced interacting partners of SARS-CoV-2 NSP6 interacting 4 proteins with miRNAs. Based on the availability of data in miRNet, we only constructed miRNA networks for the brain and lung. The degree of a node has been calculated by the number of other nodes it connected, and betweenness centrality signifies the bridging role of a node in a network. We have used the GO-BP and KEGG database for functional enrichment analysis of the miRNA network and identified significant (p ≤ 0.05) biological processes associated with the miRNA networks. Enrichment analyses were based on the hypergeometric tests after adjustment for FDR. For heart, we tried to find possible regulated pathways in a different way using miRNet. Circulating miRNAs regulated by SARS-CoV-2 infection were chosen to find out the target genes.^56^ Among these genes, only those were chosen that were regulated by SARS-CoV-2 infection in heart. miRNA-mRNA interactome for these genes and their respective miRNA were deduced.

### 2.9. Comparison of SARS-CoV-2 influenced circulatory miRNAs and tissue-specific miRNAs targeting interacting proteins of SARS-CoV-2 NSP6 interacting proteins

As SARS-CoV-2 influences miRNAs, we wanted to study the regulated circulatory miRNAs and miRNAs that target SARS-CoV-2-regulated proteins interacting with SARS-CoV-2 NSP6 binding proteins in the brain and lung. Common miRNAs in the reported list of SARS-CoV-2-influenced circulating miRNAs,^57,58^ and tissue-specific miRNAs influencing interacting proteins of SARS-CoV-2 NSP6 binding proteins were probed to find out regulated miRNAs and their respective target proteins.

## 3. Results

### 3.1. Interacting partners of NSP6 interacting proteins varies in different tissues

SARS-CoV-2 NSP6 is reported to interact with four host proteins, namely ATP synthase membrane subunit g (ATP5MG), ATPase H+ transporting accessory protein-1 (ATP6AP1), ATPase 13A3 (ATP13A3), and Sigma-1 receptor (SIGMAR1).^24^ To know the tissue-specific effect in the brain, heart, and lung, interactome analyses were done to identify interacting partners of these four NSP6 interacting proteins in each of the tissues. Proteins were selected that were regulated after COVID-19 infection^39,46,41^ as well as that were expressed in these tissues. Tissue-specific experiment-based interaction network from IID data base analyses revealed 18, 45 and 23 interacting proteins in the brain, heart and lung respectively (Supporting Information Data 1). By using Venn diagram analysis (Figure 1a), we analyzed common and unique proteins among the brain, heart, and lung. Identification of common and unique proteins would help us to identify common and unique pathways as well as probable drug targets.

**Figure 1.**
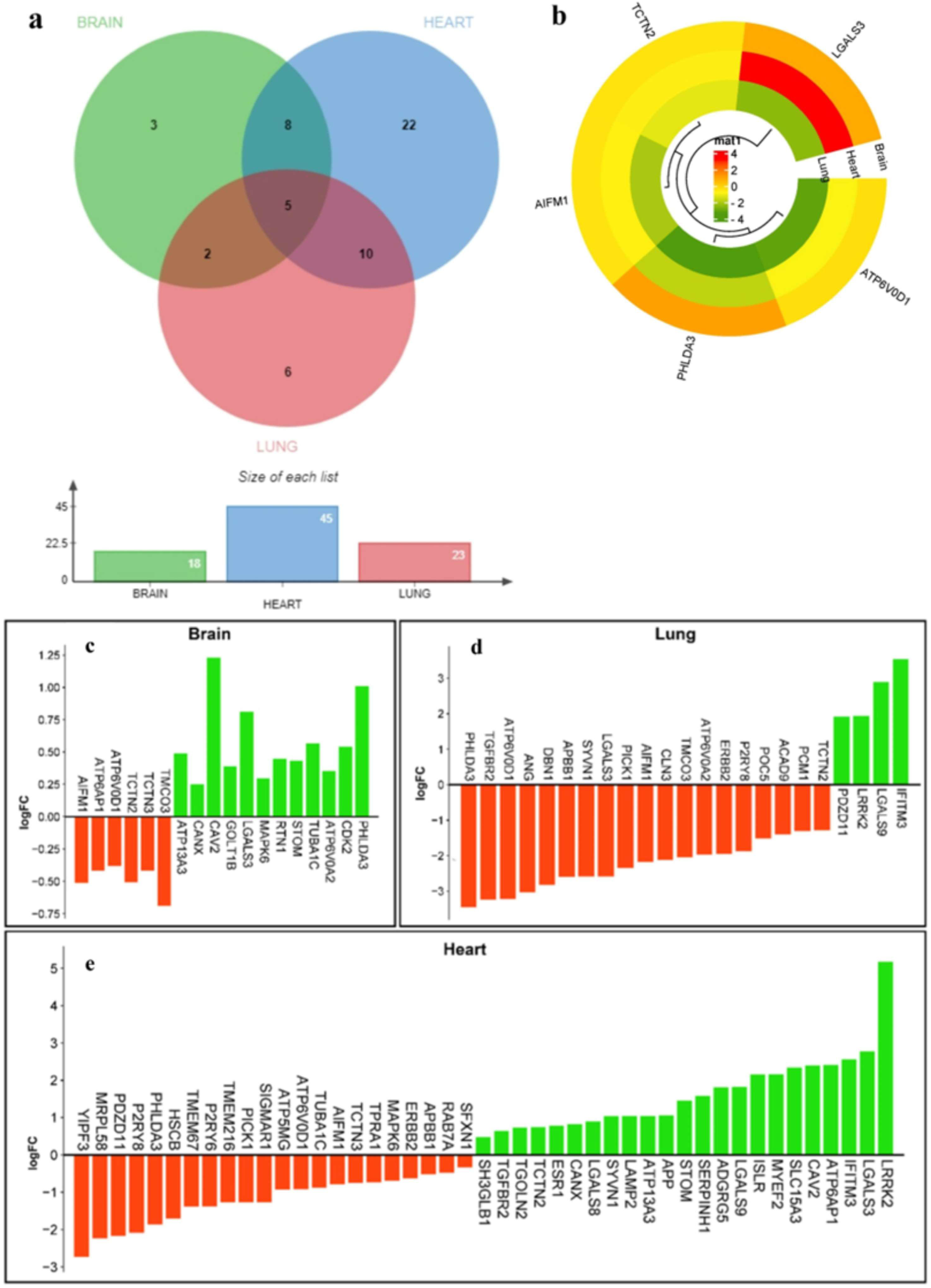
Tissue-specific COVID-19 influenced common and unique proteins interacting with SARS-CoV-2 NSP6 interacting proteins in the brain, heart, and lung. a) Venn diagram showing the number of proteins interacting with SARS-CoV-2 NSP6 interacting proteins in different tissues influenced by SARS-CoV-2 infection. Number of proteins in the brain, heart, and lung were 18, 45, and 23 respectively. Among these, 5 genes were common in the brain, lung, and heart. Common genes between the brain and heart were 8, the heart and lung were 10, and lung and the brain were 2. b) Circular cluster heatmap showing expression in fold change of the 5 common genes in the brain, heart, and lung. Regulated gene expression in the brain (c), lung (d), and the heart (e) with their fold change (Log2FC) after SARS-CoV-2 infection.

Results showed 5 of these interacting proteins were common in these three tissues and they were apoptosis inducing factor mitochondria associated 1 (AIFM1), ATPase H+ transporting V0 subunit d1 (ATP6V0D1), tectonic family member 2 (TCTN2), galectin 3 (LGALS3), and pleckstrin homology like domain family A member 3 (PHLDA3). Regulation of these genes in different tissues are shown in figure 1b. Figure 1c, 1d, and 1e shows all regulated genes in the brain, lung, and heart respectively. We further observed that ten interacting proteins interferon-induced transmembrane protein 3 (IFITM3), galectin 9 (LGALS9), P2Y receptor family member 8 (P2RY8), leucine-rich repeat kinase 2 (LRRK2), transforming growth factor beta receptor 2 (TGFBR2), amyloid beta precursor protein binding family B member (APBB1), Erb-b2 receptor tyrosine kinase 2(ERBB2), Protein interacting with PRKCA 1 (PICK1), Synoviolin 1 (SYVN1), and PDZ domain containing 11 (PDZD11) were common between the heart and lung. Two interacting proteins, transmembrane and coiled-coil domains 3 (TMCO3), and ATPase H+ transporting V0 subunit a2 (ATP6V0A2) were common in the brain and lung, while in the brain and heart, eight shared proteins were observed, namely, ATPase H+ transporting accessory protein 1 (ATP6AP1), tectonic family member 3 (TCTN3), ATPase 13A3 (ATP13A3), calnexin (CANX), caveolin 2 (CAV2), mitogen-activated protein kinase 6 (MAPK6), stomatin (STOM), and tubulin alpha 1c (TUBA1C).

Considering tissue-specific regulated unique proteins, only three were present in the brain namely golgi transport 1B (GOLT1B), reticulon 1 (RTN1), cyclin-dependent kinase 2 (CDK2). 22 proteins were unique in the heart and they were Yip1 domain family member 3 (YIPF3), mitochondrial ribosomal protein L58 (MRPL58), amyloid beta precursor protein (APP), immunoglobulin superfamily containing leucine-rich repeat (ISLR), serpin family H member 1 (SERPINH1), lysosomal-associated membrane protein 2 (LAMP2), transmembrane protein 216 (TMEM216), adhesion G protein-coupled receptor G5 (ADGRG5), ATP synthase membrane subunit g (ATP5MG), trans-golgi network protein 2 (TGOLN2), HscB mitochondrial iron-sulfur cluster co-chaperone (HSCB), transmembrane protein 67 (TMEM67), myelin expression factor 2 (MYEF2), member RAS oncogene family (RAB7A), pyrimindinergic receptor P2Y6 (P2RY6), solute carrier family 15 member 3 (SLC15A3), sigma non-opioid intracellular receptor 1 (SIGMAR1), transmembrane protein adipocyte associated 1 (TPRA1), SH3 domain-containing GRB2-like (SH3GLB1), estrogen receptor 1 (ESR1), galectin 8 (LGALS8), sideroflexin 1 (SFXN1). In the lungs, six unique proteins observed were angiogenin (ANG), drebrin 1 (DBN1), battenin(CLN3), POC5 centriolar protein (POC5), acyl-CoA dehydrogenase family member 9 (ACAD9), and pericentriolar material 1 (PCM1).

### 3.2. Hub genes network and its associated pathways

#### 3.2.1. Hub genes network

After finding out various proteins interacting with SARS-CoV-2 NSP6 in a tissue-specific manner, we wanted to find out networks and the prominent players. For this STRING database was used. In the STRING database, the compilation of tissue-specific regulated genes pertaining to the brain, heart, and lung was employed to construct a Protein-Protein (PP) Interactome network, consisting of the query proteins and their primary interactors of the first shell. In the context of the brain tissue, this interactome yielded a network characterized by the following features: a total count of 28 nodes, accompanied by 46 edges, and thereby resulting in an average node degree of 3.29. Similarly in the heart, the PP interactome revealed 50 nodes corresponding to 70 edges along with an average node degree of 2.8. Whereas in the lung, the PP interactome consisted of 33 nodes and 38 edges, having an average node degree of 2.3.

For the key genes in the PP interactome network of the brain, lung, and heart, we identified top 10 hub genes in cytoHubba (Supporting Information Data 2). In the brain, these hub genes were cyclin D1 (CCND1), cyclin-dependent kinase 2 (CDK2), cyclin A1 (CCNA1), cell division cycle 6 (CDC6), cyclin-dependent kinase inhibitor 1B (CDKN1B), S-phase kinase-associated protein 1 (SKP1), cyclin B1 (CCNB1), cyclin E1 (CCNE1), ATPase H+ transporting V1 subunit B1 (ATP6V1B1), and ATPase H+ transporting V1 subunit E1 (ATP6V1E1). Among these, CCND1 was the most significant one (Figure 2a). Similarly in the heart, the ras-related protein Rab-7a (RAB7A), ATPase H+ transporting V0 subunit d1 (ATP6V0D1), lysosomal-associated membrane protein 2 (LAMP2), transforming growth factor beta 1 (TGFB1), calnexin (CANX), galectin 3 (LGALS3), ATPase H+ transporting V1 subunit subunit B1 (ATP6V1B1), ATPase H+ transporting V1 subunit D (ATP6V1D), ATPase H^+^-transporting accessory protein 1 (ATP6AP1), and Amyloid beta precursor protein (APP) were the hub genes. RAB7A, ATP6V0D1, and LAMP2 were more significant than other hub genes (Figure 2b). In the case of the lungs, ATPase H+ transporting V0 subunit a1 (ATP6V0A1), ATPase H+ transporting V0 subunit a2 (ATP6V0A2), ATP6V0D1, ATP6V1D, ATPase H+ transporting V1 subunit E1 (ATP6V1E1), Transforming growth factor beta 1 (TGFB1), Epidermal growth factor receptor-binding protein 1 (EGFB1), Epidermal growth factor (EGF), Transforming growth factor beta receptor 2 (TGFBR2), and Galectin 3 (LGALS3) were identified as hub genes (Figure 2c).

**Figure 2a.**
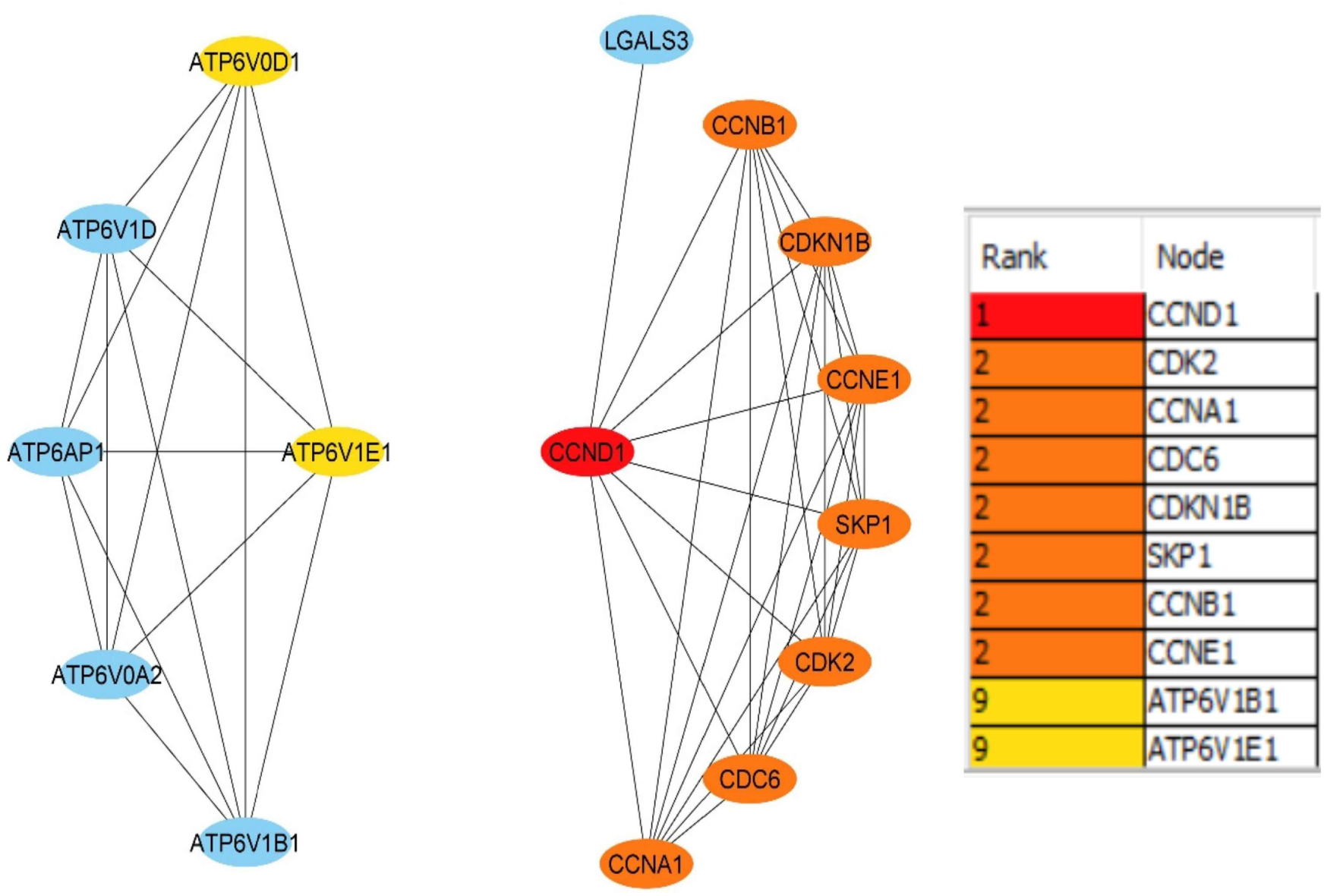
Top 10 hub genes of the NSP6 interactome proteins from STRINGdb analyzed by cytoHubba in the brain. Node color from red to yellow represents higher to lower significance. Analysis showed CCND1 to be the highly significant major hub gene.

**Figure 2b.**
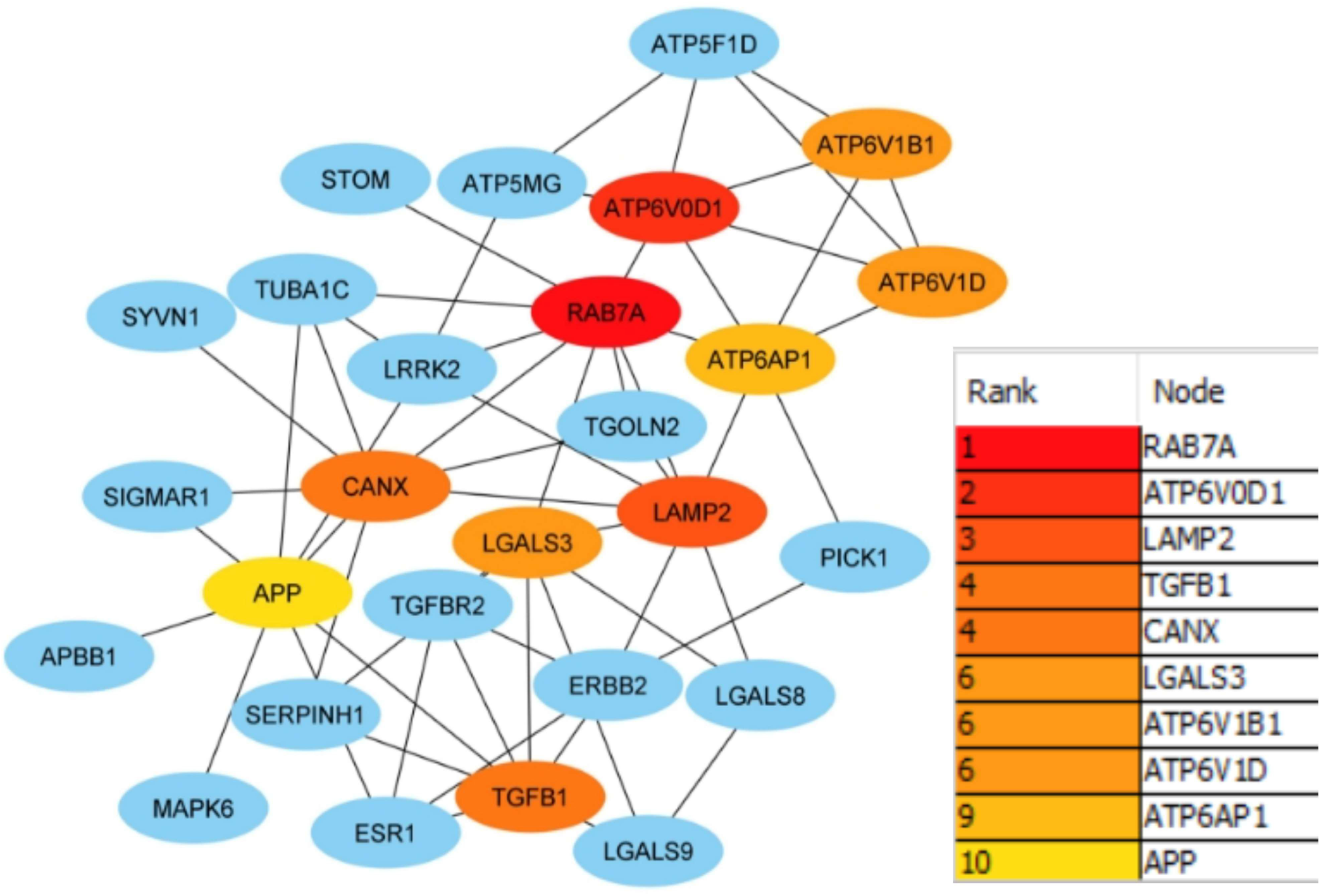
Top 10 hub genes of the NSP6 interactome proteins from STRINGdb analyzed by cytoHubba in the heart. Node color from red to yellow represents higher to lower significance. Analysis showed RAB7A, ATP6V0D1, and LAMP2 to be the highly significant major hub genes.

**Figure 2c.**
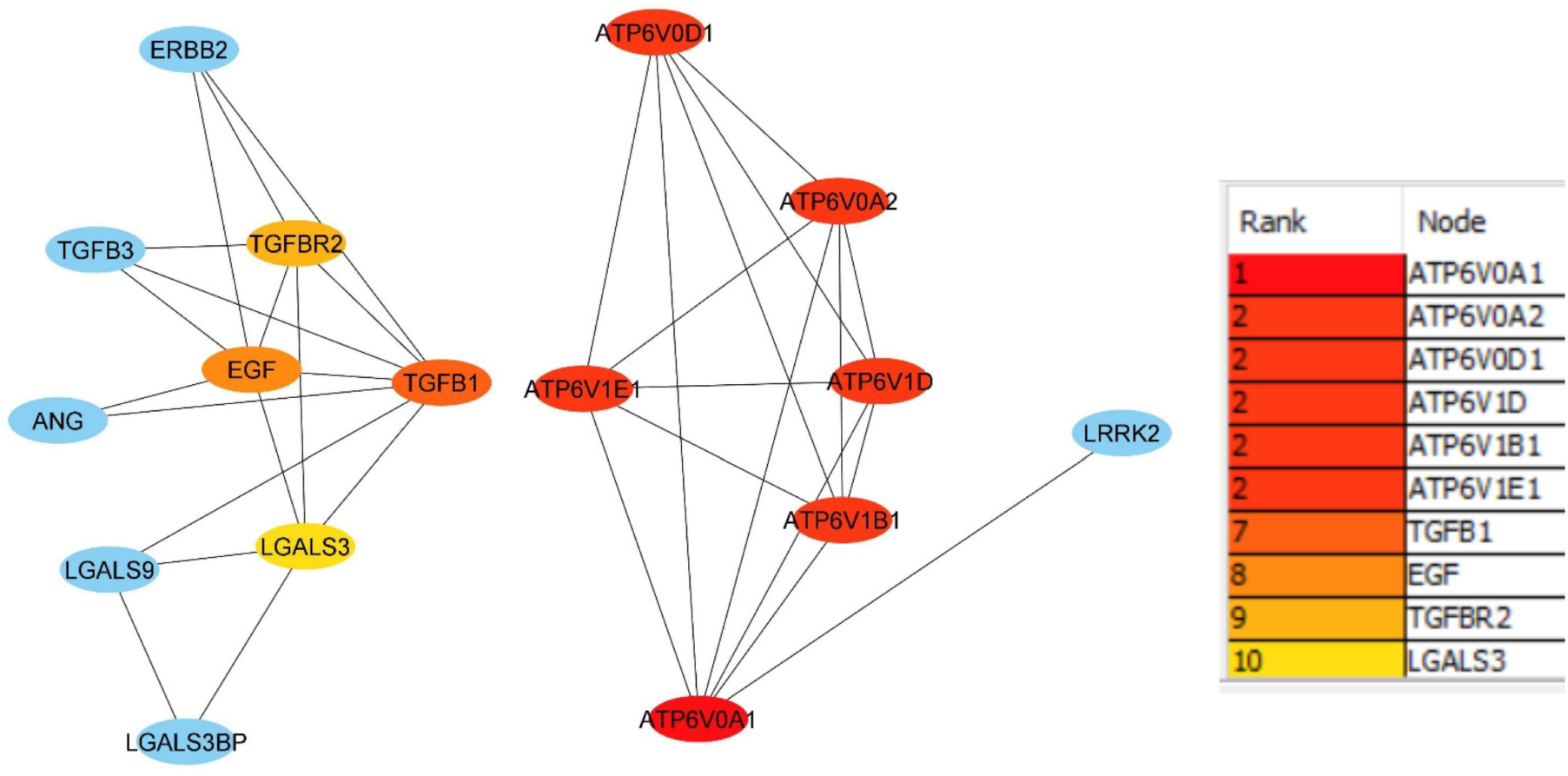
Top 10 hub genes of the NSP6 interactome proteins from STRINGdb analyzed by cytoHubba in the lung. Node color from red to yellow represents higher to lower significance. Analysis showed ATP6V0A1, ATP6V0A2, ATP6V0D1, ATP6V1D, ATP6V1Bl, and ATP6V1El to be the highly significant major hub genes.

#### 3.2.2. Hub genes associated pathways

Pathway enrichment analyses associated with the top 10 hub genes from the brain, lung, and heart were performed to identify the regulated biological and cellular processes. In case of the brain, iron uptake and transport, and G1/S transition were the two main key biological ontologies found. In addition, 97% of biological pathways were associated with G1/S transition showing interaction with 8 of the total hub genes. In addition, two pathways, namely small cell lung cancer (CDK2, CDKN1B, CCNE1, CCND1), and prostate cancer (CDK2, CDKN1B, CCNE1, CCND1) were also identified. Also activation of PTK6 and regulation of cell cycle associated pathways, and p53 signalling pathways were also found in the analysis (Figure 3a).

**Figure 3a.**
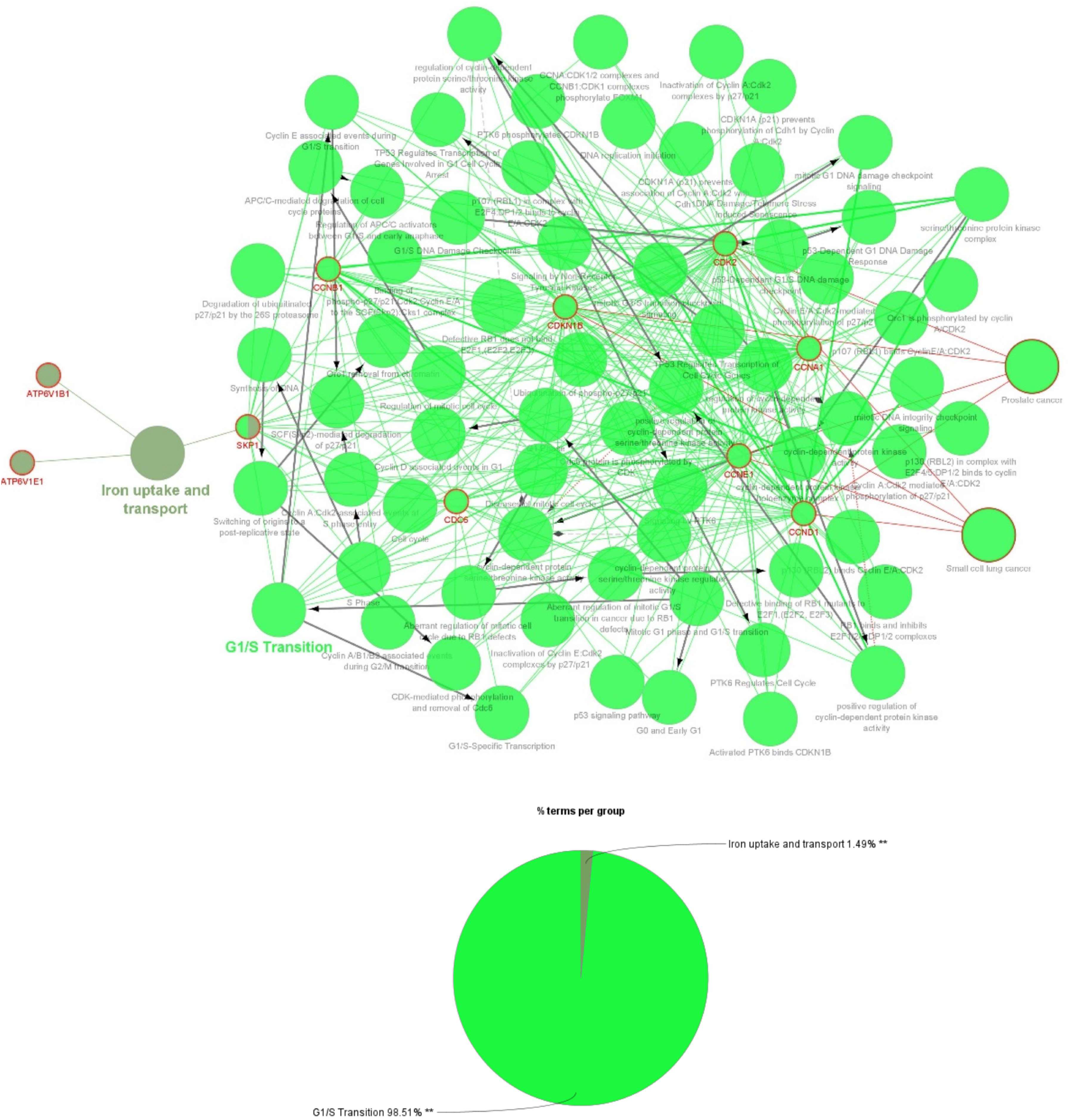
Gene ontology processes, KEGG pathway and reactome pathway analyses of top 10 hub genes in the brain by CluGO analysis.

In the context of lung biology, two prominent biological pathways had been identified as crucial and they were the differentiation of dendritic cells and the reduction of intraphagosomal pH to a level of 5, primarily facilitated by V-ATPase. Furthermore, it had been observed that 97.22% of biological pathways were linked to the reduction of intraphagosomal pH, indicating significant interaction with six of the total hub genes. Notably, our analyses also revealed associations with other processes, including rheumatoid arthritis, insulin receptor cycling, V*ibrio cholerae* infection, and epithelial cell signaling in *Helicobacter pylori* infection. Among these hub genes, LGALS3, TGFBR2, and TGFB1 were specifically associated with the dendritic cell differentiation pathway (Figure 3b).

**Figure 3b.**
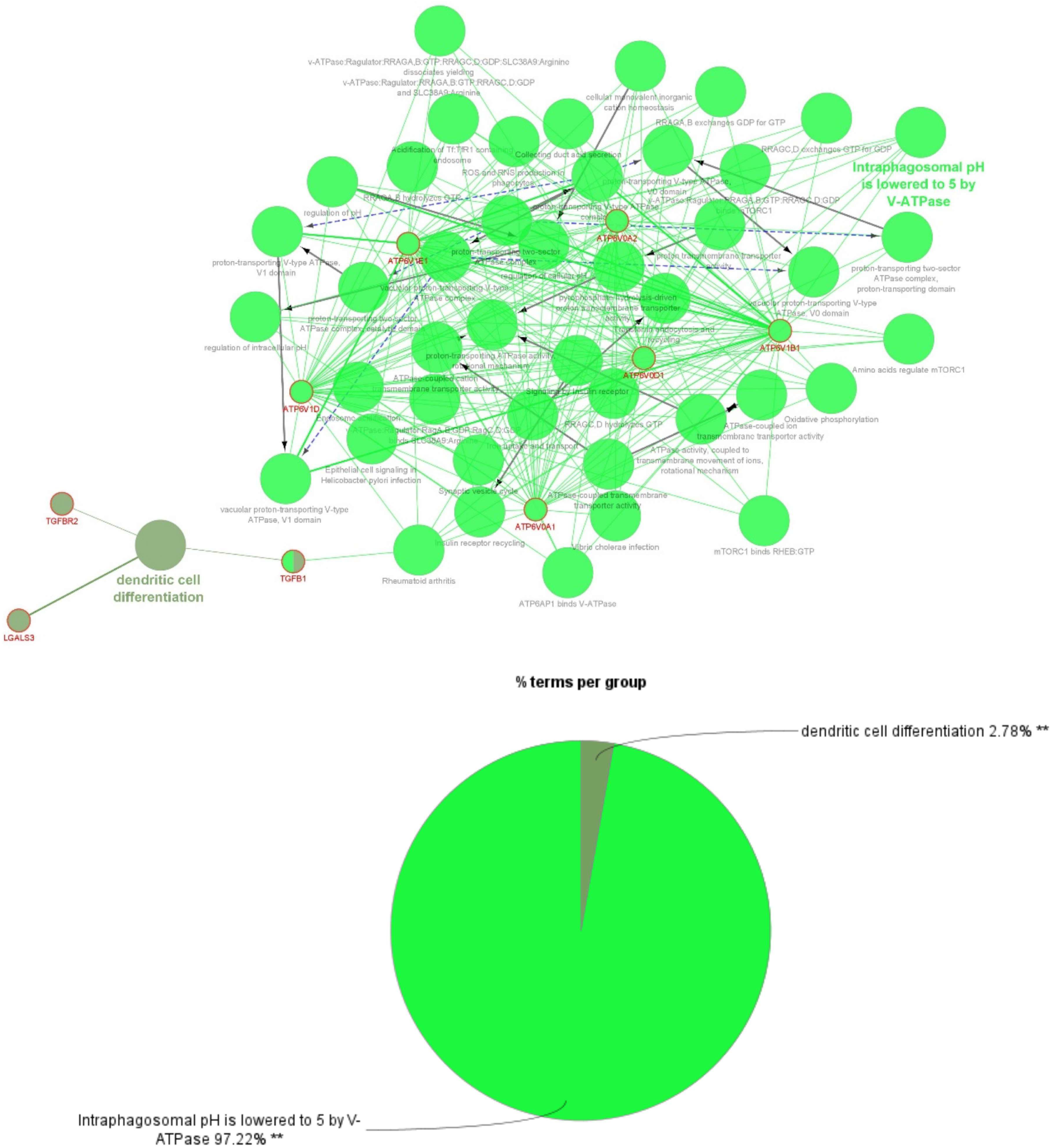
Gene ontology processes, KEGG pathway and reactome pathway analyses of top 10 hub genes in the lung by CluGO analysis.

For the heart ClueGO analysis revealed, phagosome associated pathways (2.86% terms per group), and regulation of intracellular pH (97.14% terms per group) as the leading terms of the network. Proteins associated with phagosome associated pathways had close interaction with all the hub genes (ATP6V1B1, ATP6V1D, ATP6AP1, ATP6V0D1, RAB7A, CANX, and LAMP2) except TGFB1. Proteins associated with regulation of intracellular pH were ATP6V1B1, ATP6V1D, ATP6AP1, ATP6V0D1, and RAB7A. Some of the functions influenced by the genes of the network were vascular proton-transporting V-type ATPase complex (ATP6V1B1, ATP6V1D, and ATP6V0D1), rheumatoid arthritis (TGFB1, ATP6V1B1, ATP6V1D, ATP6AP1, and ATP6V0D1), and additional pathways like mTORC1 pathways, iron uptake and transport, signalling by insulin receptor, ATPase-coupled cation transmembrane transporter activity etc. (Figure 3c).

**Figure 3c.**
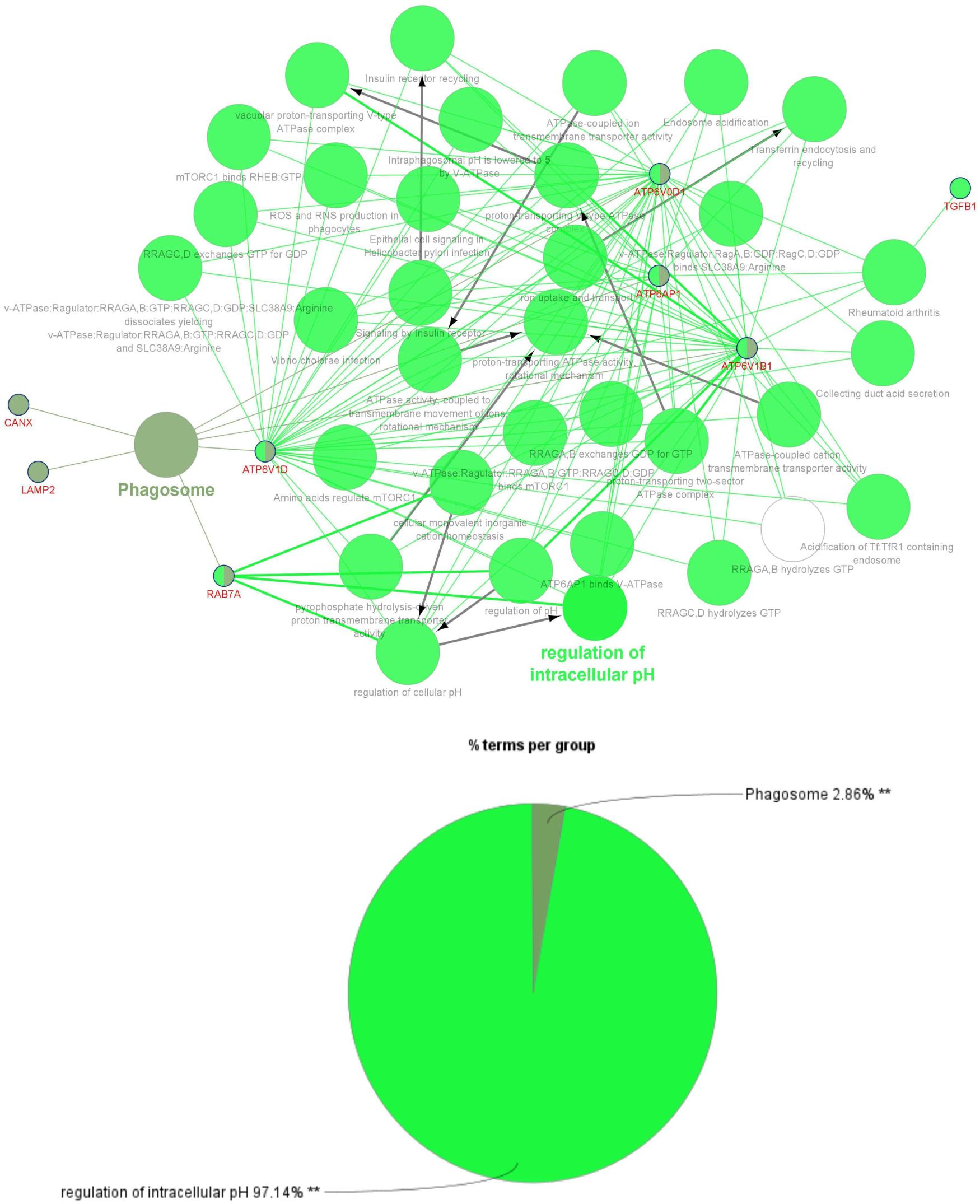
Gene ontology processes, KEGG pathway and reactome pathway analyses of top 10 hub genes in the heart by CluGO analysis.

### 3.3. Drug-gene interaction analysis of hub genes in DGIdb

After deducing pathways where the hub genes were involved, we employed DGIdb to assess potential drug interactions with the hub genes in the brain, heart, and lung. In the brain among the top ten hub genes, five genes (CDKN1B, CCND1, CDK2, CCNB1, and CCNE1) were drug targets. Few drugs were targeting multiple proteins. There were 30 approved drugs, and 89 non-approved drugs (Figure 4a) (Supporting Information Data 3). In the heart (TGFB1, CANX, APP, and LGALS3) and in the lungs (TGFB1, LGALS3, TGFBR2, and EGF), four genes each were drug targets (Figure 4b and 4c respectively and Supporting Information Data 3). Approved and non-approved drugs in the heart were 44 and 56 respectively, and in the lung, the counts were 34 and 31 respectively.

**Figure 4a.**
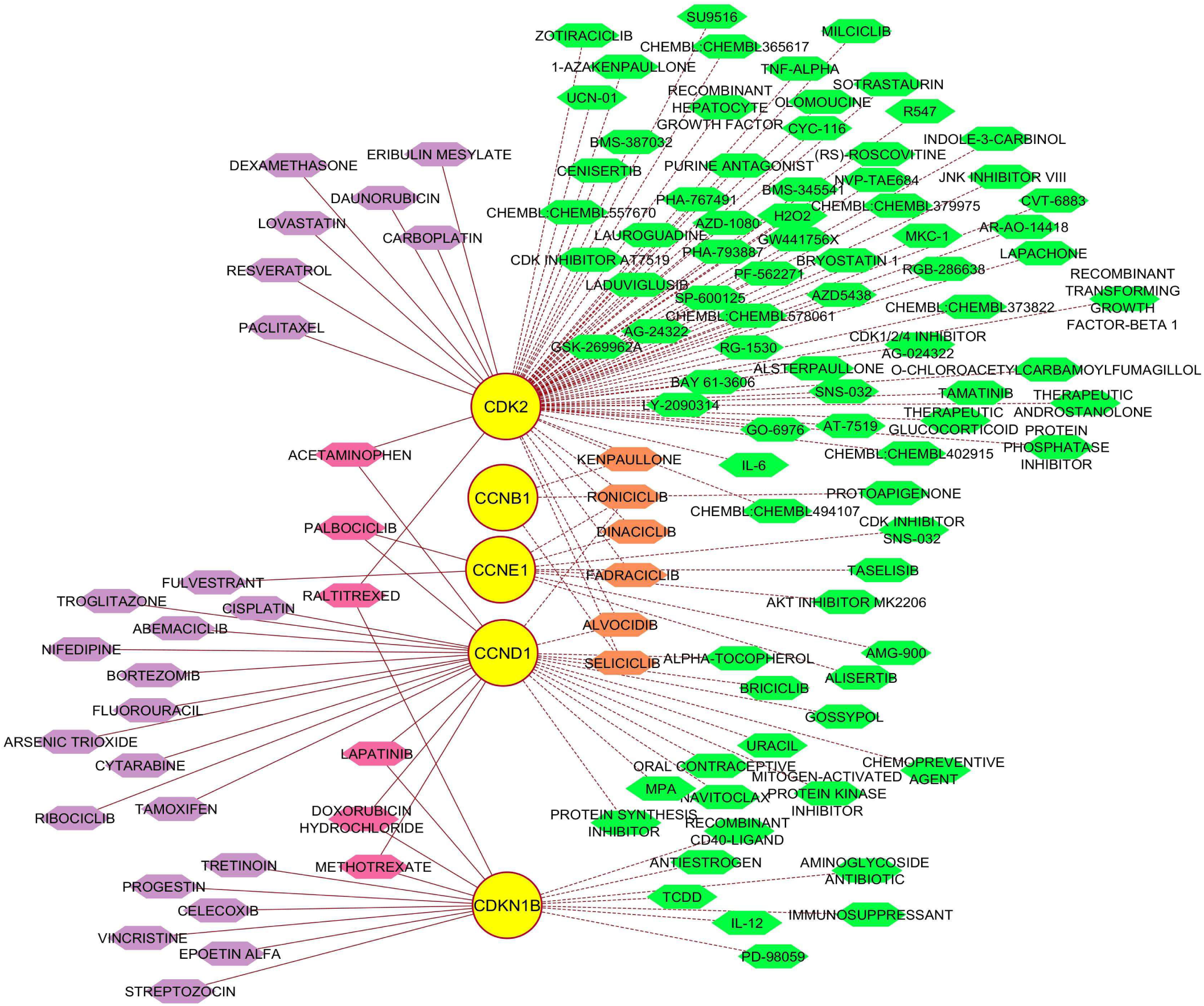
Drug targets among hub genes in the brain. Circular yellow nodes are hub genes and hexagonal nodes denote drugs. Violet hexagonal nodes are approved drugs and pink nodes denote approved drugs targeting more than one hub gene. Green colored hexagonal nodes with dotted lines are non-approved drugs. Solid edges are approved drugs and dotted edges are for non-approved drugs. Orange colored hexagonal nodes denote non-approved drugs targeting more than one hub gene. 30 approved and 89 non-approved drugs were found for the five among 10 top hub genes.

**Figure 4b.**
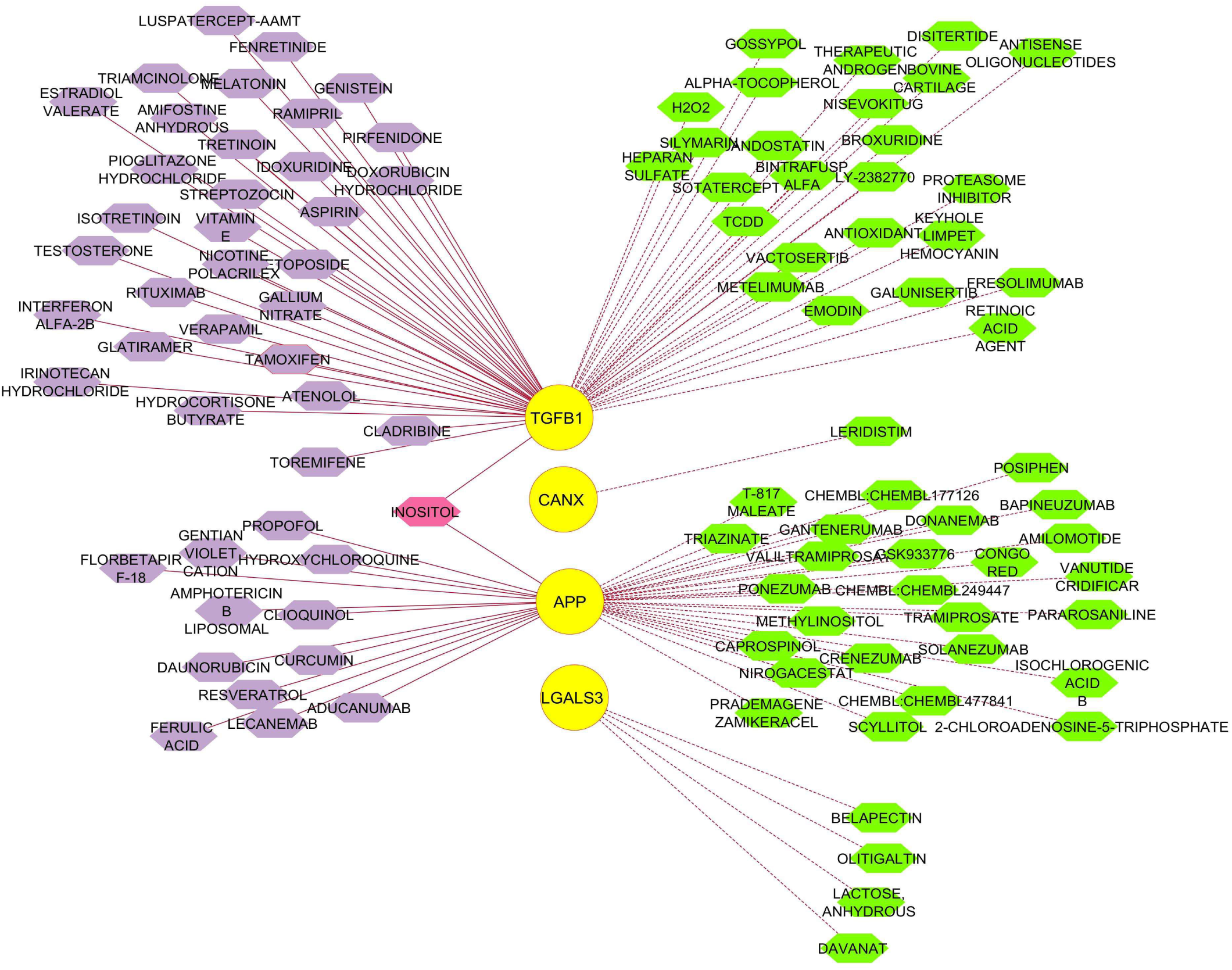
Drug targets among hub genes in the heart. Circular yellow nodes are hub genes and hexagonal nodes denote drugs. Violet hexagonal nodes are approved drugs and pink nodes denote approved drugs targeting more than one hub gene. Solid edges are approved drugs and dotted edges are for non-approved drugs. Green colored hexagonal nodes are non-approved drugs. 44 approved and 56 non-approved drugs were found for the four among 10 top hub genes.

**Figure 4c.**
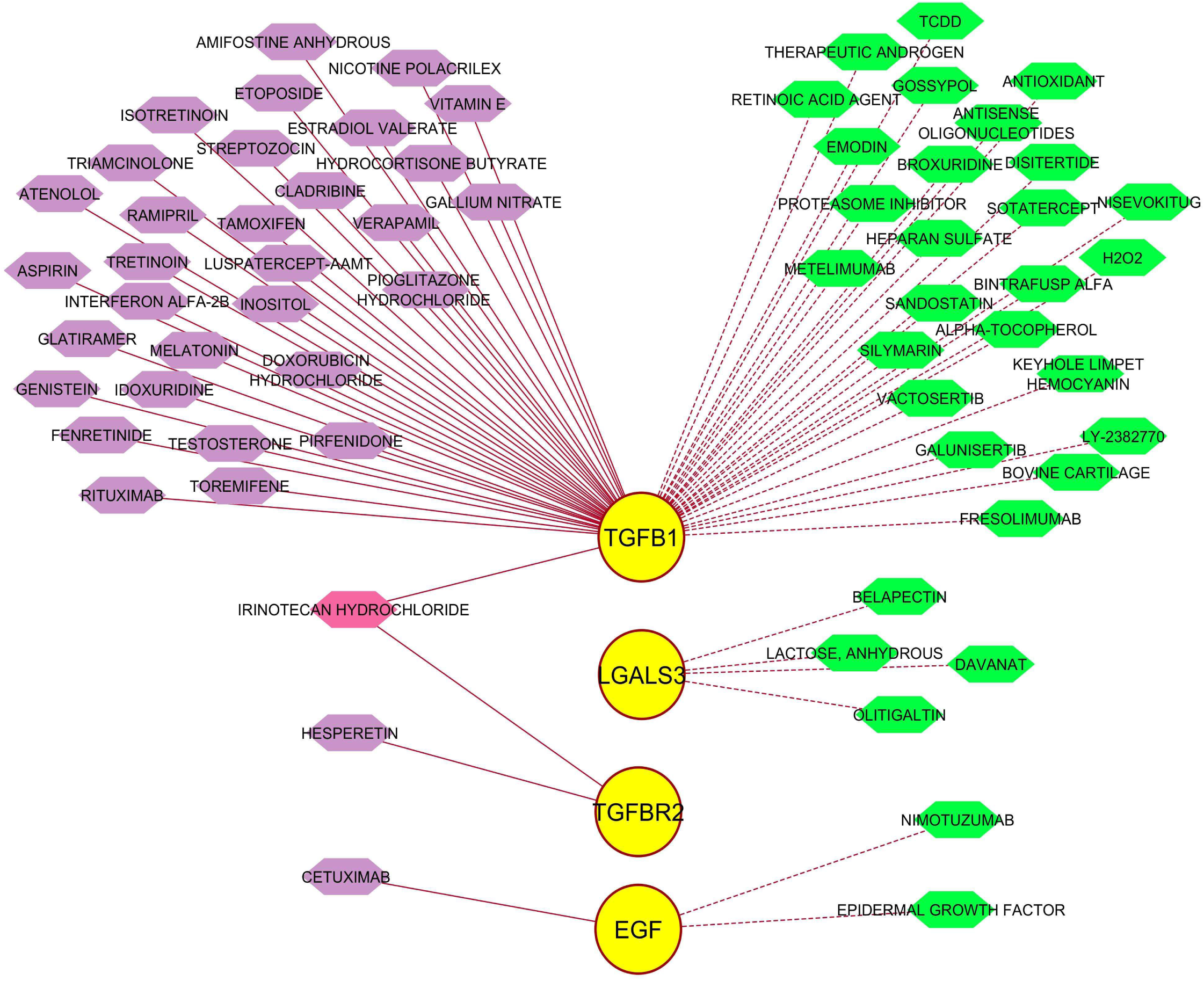
Drug targets among hub genes in the lung. Circular yellow nodes are hub genes and hexagonal nodes denote drugs. Violet hexagonal nodes are approved drugs and pink nodes denote approved drugs targeting more than one hub gene. Solid edges are approved drugs and dotted edges are for non-approved drugs. Green colored hexagonal nodes are non-approved drugs. 34 approved and 31 non-approved drugs were found for the four among 10 top hub genes.

### 3.4. NSP6 interacting proteins can influence general and unique tissue-specific biological processes

Here, tissue-specific interacting partners of NSP6 interacting proteins had been used to bring out the related pathways those are the most susceptible and probably influenced due to COVID-19 infections in the brain, heart, and lungs. Pathways in the brain, heart, and lungs were 25, 147, and 168 respectively. There were 2, 52, and 5 common GO pathways between brain-lung, lung-heart, and heart-brain pairs respectively and 14 common GO pathways were present in all. This aided in identifying pathways of notable significance across all of these tissue types. Gene Ontology (GO) enrichment analysis revealed 4, 69, and 100 unique pathways that were present in the brain, heart, and lung respectively (Figure 5 and Supporting Information Data 4). Common pathways in the brain, heart, and lung are shown in figure 6 and supporting information data 4. It is observed that for the same pathway in different organs the genes involved may differ.

**Figure 5.**
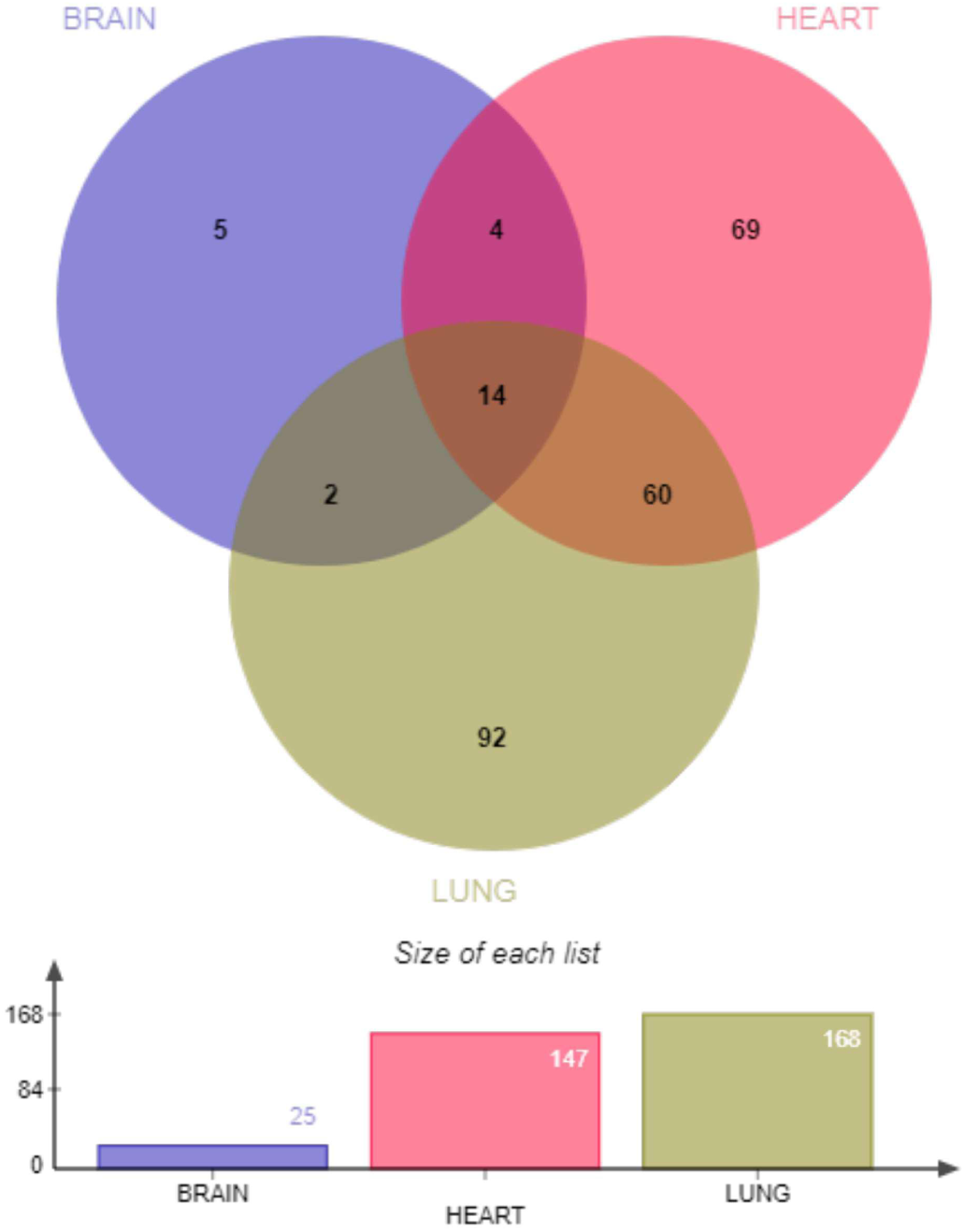
Venn diagram representing common and unique pathways influenced by proteins interacting with SARS-CoV-2 NSP6 interacting proteins in the brain, heart, and lungs. 14 pathways were common in all the three tissues, whereas unique pathways present in the brain, heart, and lungs were 4, 69, and 100 respectively.

**Figure 6.**
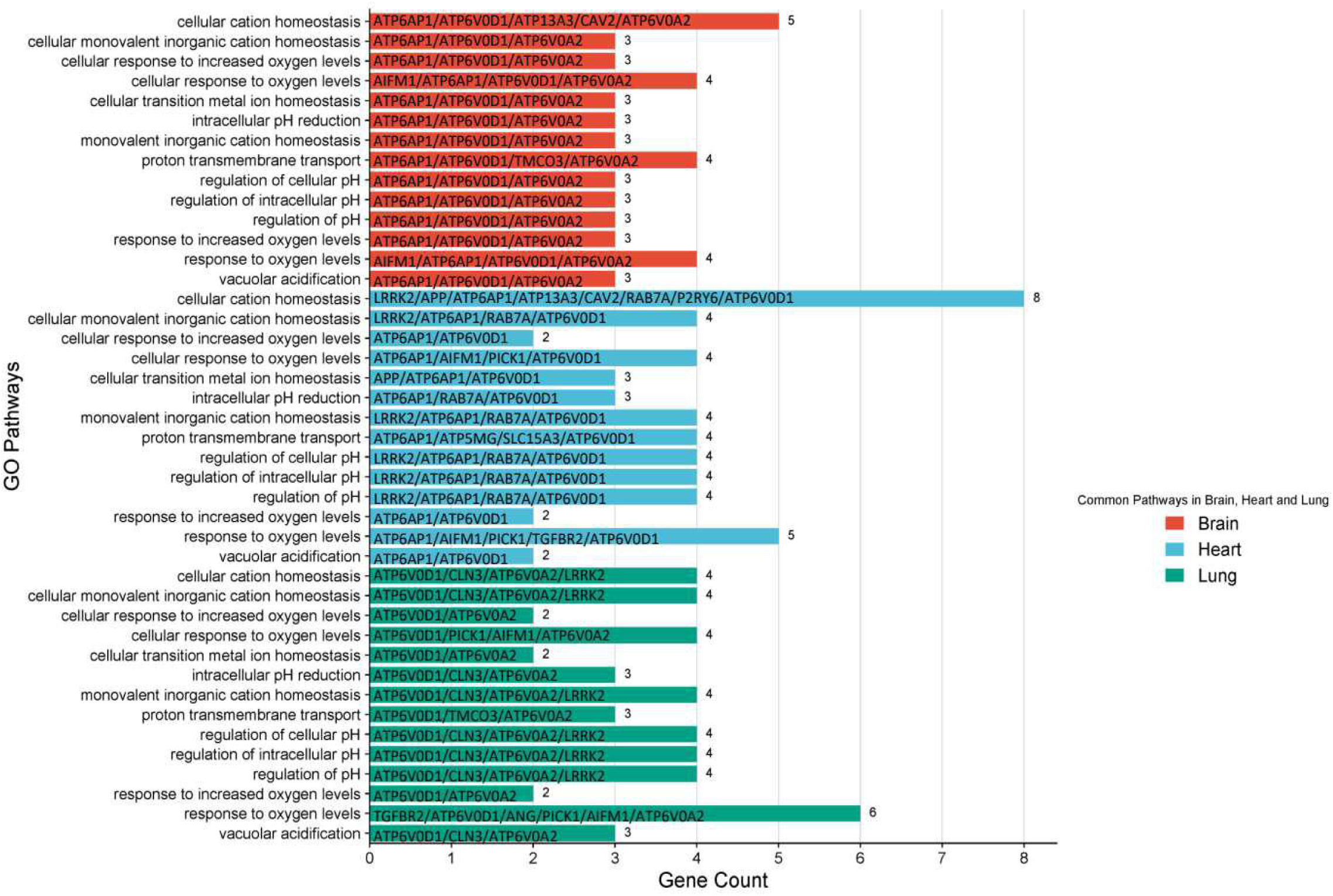
Bar diagram showing common pathways influenced by proteins interacting with SARS-CoV-2 NSP6 interacting proteins in the brain, heart, and lungs. Length of the bars represents the number of genes involved in the pathways and the genes involved are mentioned inside the bars. For the same pathway in different organs the set of genes involved may differ. X-axis represents gene count and Y-axis represents GO terms.

Generally, SARS-CoV-2 NSP6 triggers autophagy activity via the omegasome pathway,^59^ inducing the ER zippering activity,^29^ and inducing inflammasome activation in lung epithelial cells.^60^ In addition, among these three major organs (the brain, heart, and lung), NSP6 influenced pathways were noticed to be more in the heart and lung than in the brain. In the brain, unique pathways included response to nitric oxide, golgi lumen acidification, cellular response to reactive nitrogen species, cellular response to nitric oxide, and modulation by host of symbiont processes (Figure 7a and Supporting Information Data 4).

**Figure 7a.**
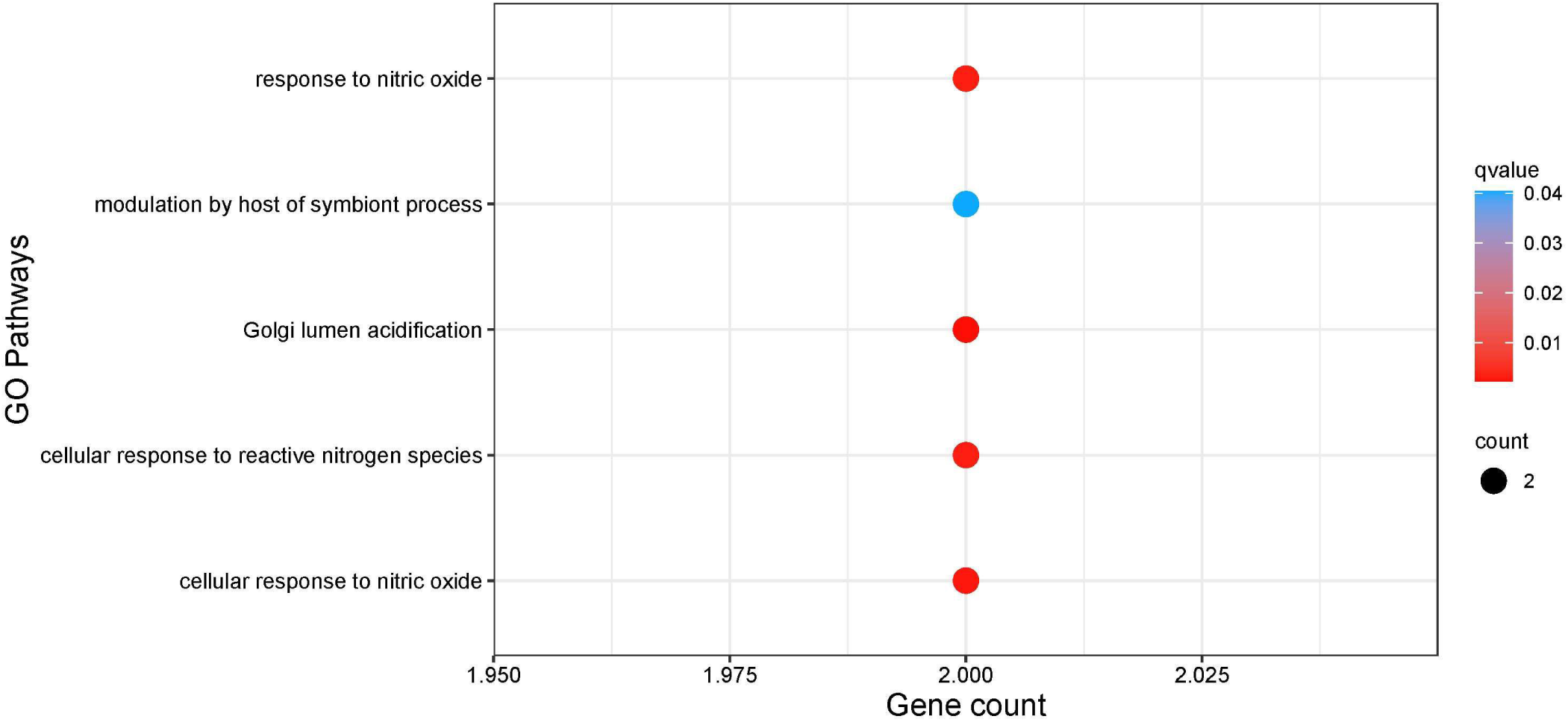
Dot plot representing unique pathways influenced by proteins interacting with SARS-CoV-2 NSP6 interacting proteins in the brain. Dot size represents the gene ratio and dot color depicts the q-value as in the heat map. X-axis shows the count of genes per GO term and Y-axis indicates the GO terms. The color gradient indicates the q-value using the Benjamini-Hochberg method.

In the heart, NSP6 can also influence some unique biological pathways including viral process, regulation of viral process, viral life cycle, biological process involved in symbiotic interaction, cellular response to starvation, regulation of dopamine receptor signaling pathway, G protein-coupled purinergic nucleotide receptor signaling pathway, endosome to plasma membrane protein transport, chondrocyte differentiation involved in endochondral bone morphogenesis, cellular response to manganese ion, etc., (Figure 7b and Supporting Information Data 4).

**Figure 7b.**
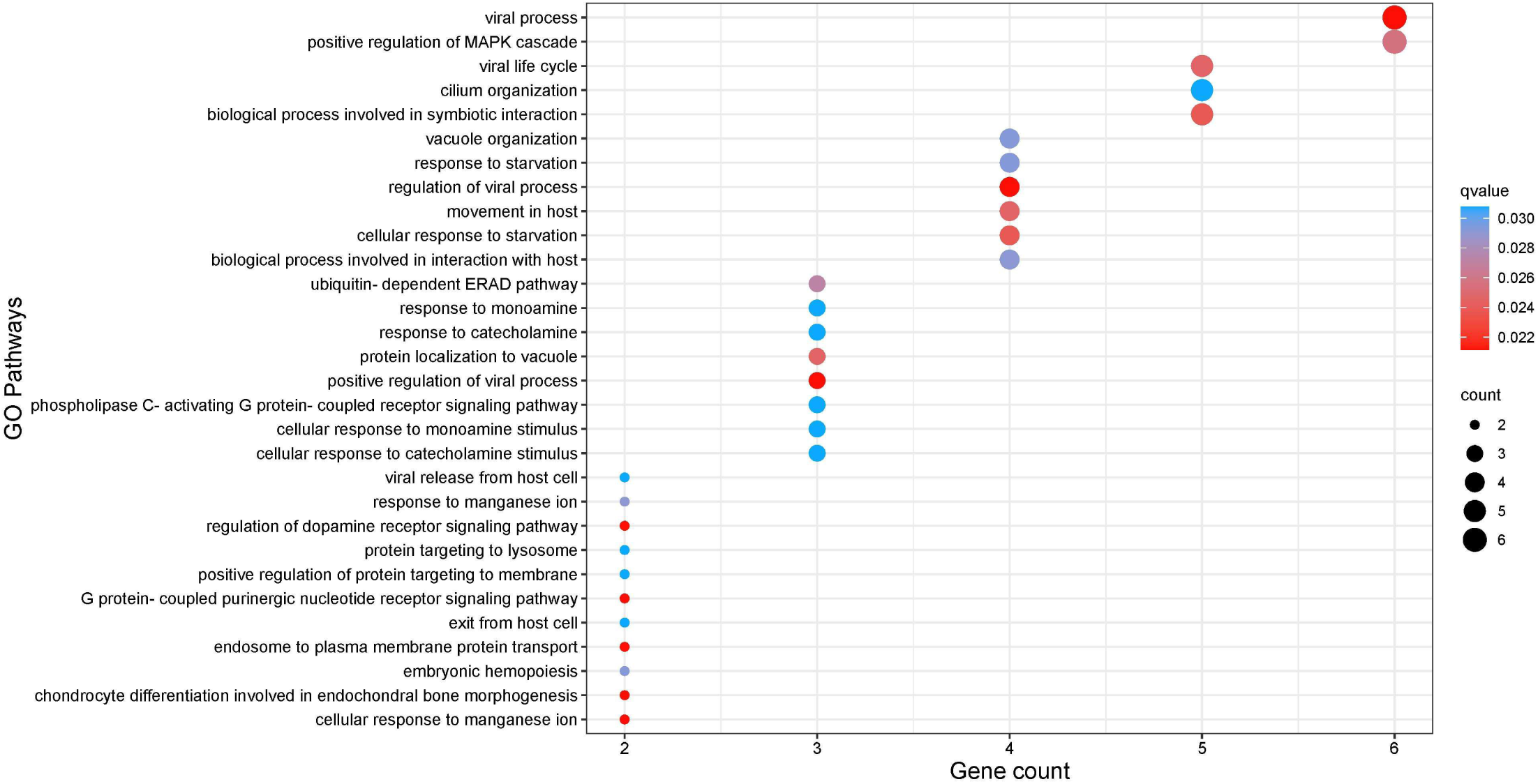
Dot plot representing top 30 unique pathways influenced by proteins interacting with SARS-CoV-2 NSP6 interacting proteins in the heart. Dot size represents the gene ratio and dot color depicts the q-value as in the heat map. X-axis shows the count of genes per GO term and Y-axis indicates the GO terms. The color gradient indicates the q-value using the Benjamini-Hochberg method.

In the case of the lung, synapse organisation, regulation of endocytosis, dendritic spine organization, response to decreased oxygen levels, regulation of T-cell activation and apoptotic process, regulation of leukocyte proliferation and leukocyte cell-cell adhesion, lymphocyte proliferation, and intrinsic apoptotic cell signalling were prominent (Figure 7c and Supporting Information Data 4).

**Figure 7c.**
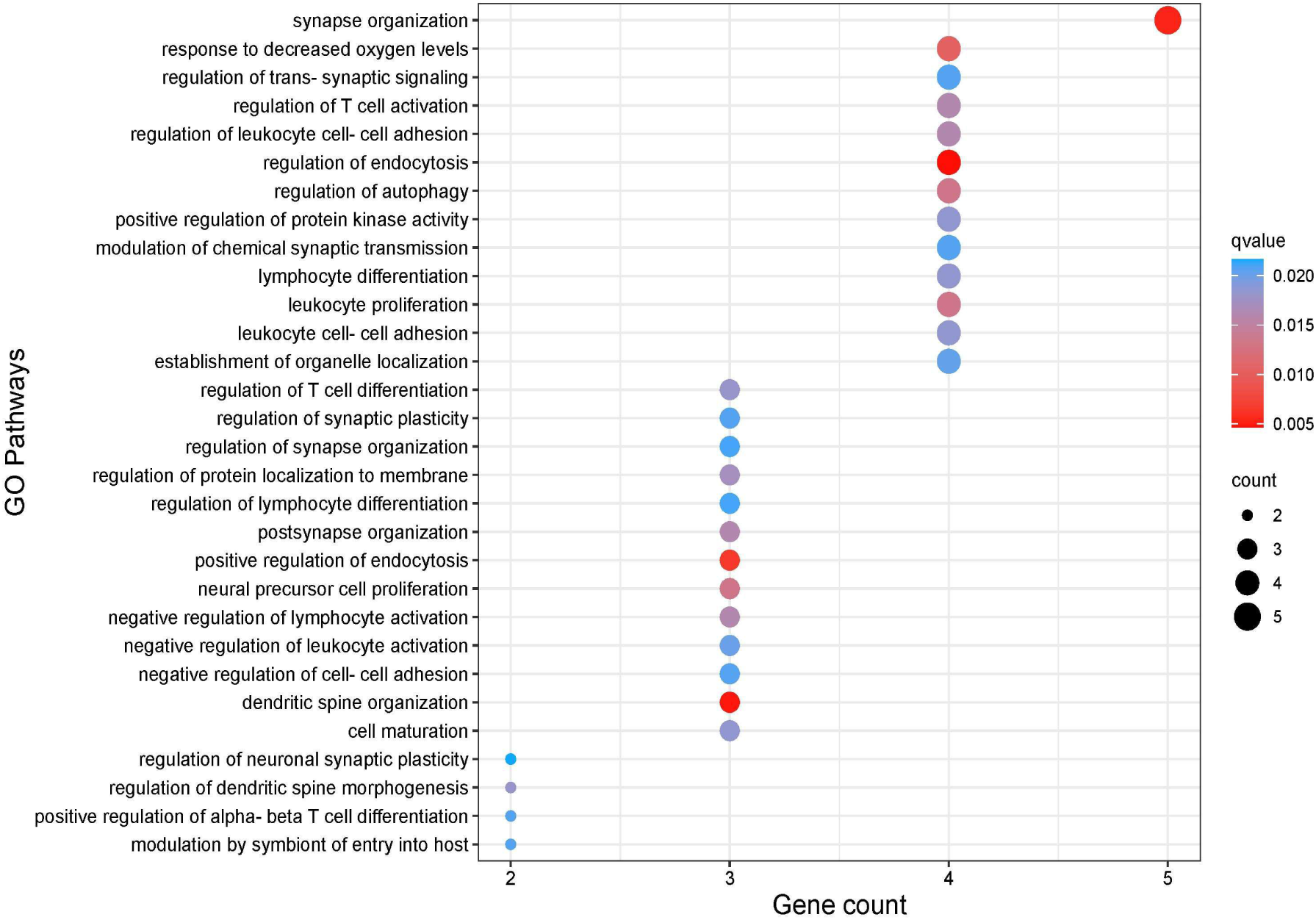
Dot plot representing top 30 unique pathways influenced by proteins interacting with SARS-CoV-2 NSP6 interacting proteins in the lungs. Dot size represents the gene ratio and dot color depicts the q-value as in the heat map. X-axis shows the count of genes per GO term and Y-axis indicates the GO terms. The color gradient indicates the q-value using the Benjamini-Hochberg method.

In the subsequent phase of our study, taking into account of the tissue-specific context, we aimed to conduct an analysis of pathways impacted by both upregulated and downregulated genes, after SARS-CoV-2 infection. We delineated both up and down regulated genes to identify pathways they influence. In the brain, among the upregulated genes impacted pathways, the prominent ones were positive regulation, and modulation of host viral process (Figure 8a and Supporting Information Data 5).

**Figure 8a.**
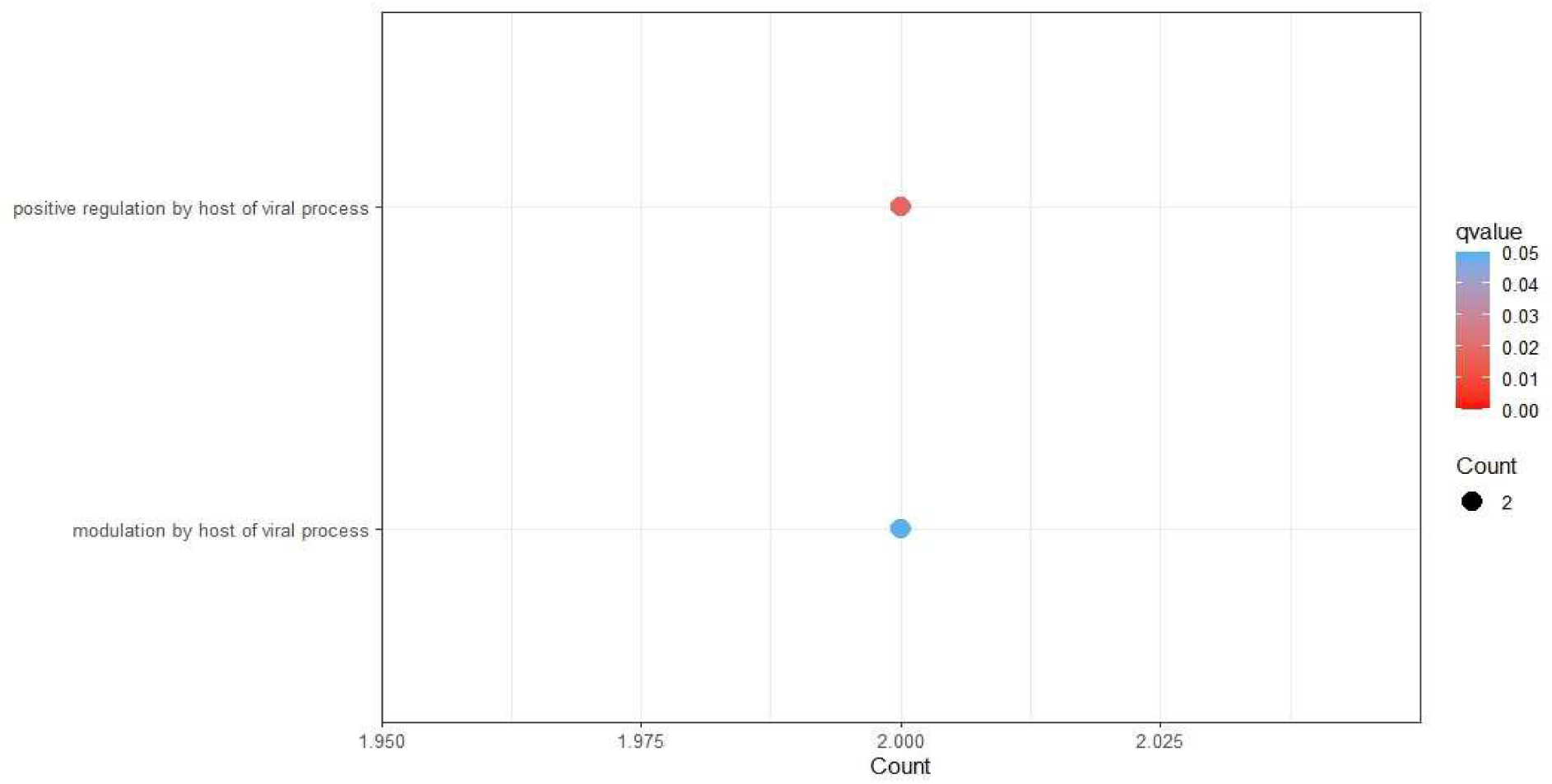
Dot plot representing pathways in the brain influenced by upregulated proteins interacting with SARS-CoV-2 NSP6 interacting proteins. Dot size represents the gene ratio and dot color depicts the q-value as in the heat map. X-axis shows the count of genes per GO term and Y-axis indicates the GO terms. The color gradient indicates the q-value using the Benjamini-Hochberg method.

The downregulated genes were observed to be prominantly associated with cellular response to oxygen levels, proton transmembrane transport, cilium organization and assembly, regulation of cellular and intracellular pH, vacuolar acidification, transission metal ion homeostasis, iron ion homeostasis, etc. (Figure 8b and Supporting Information Data 5).

**Figure 8b.**
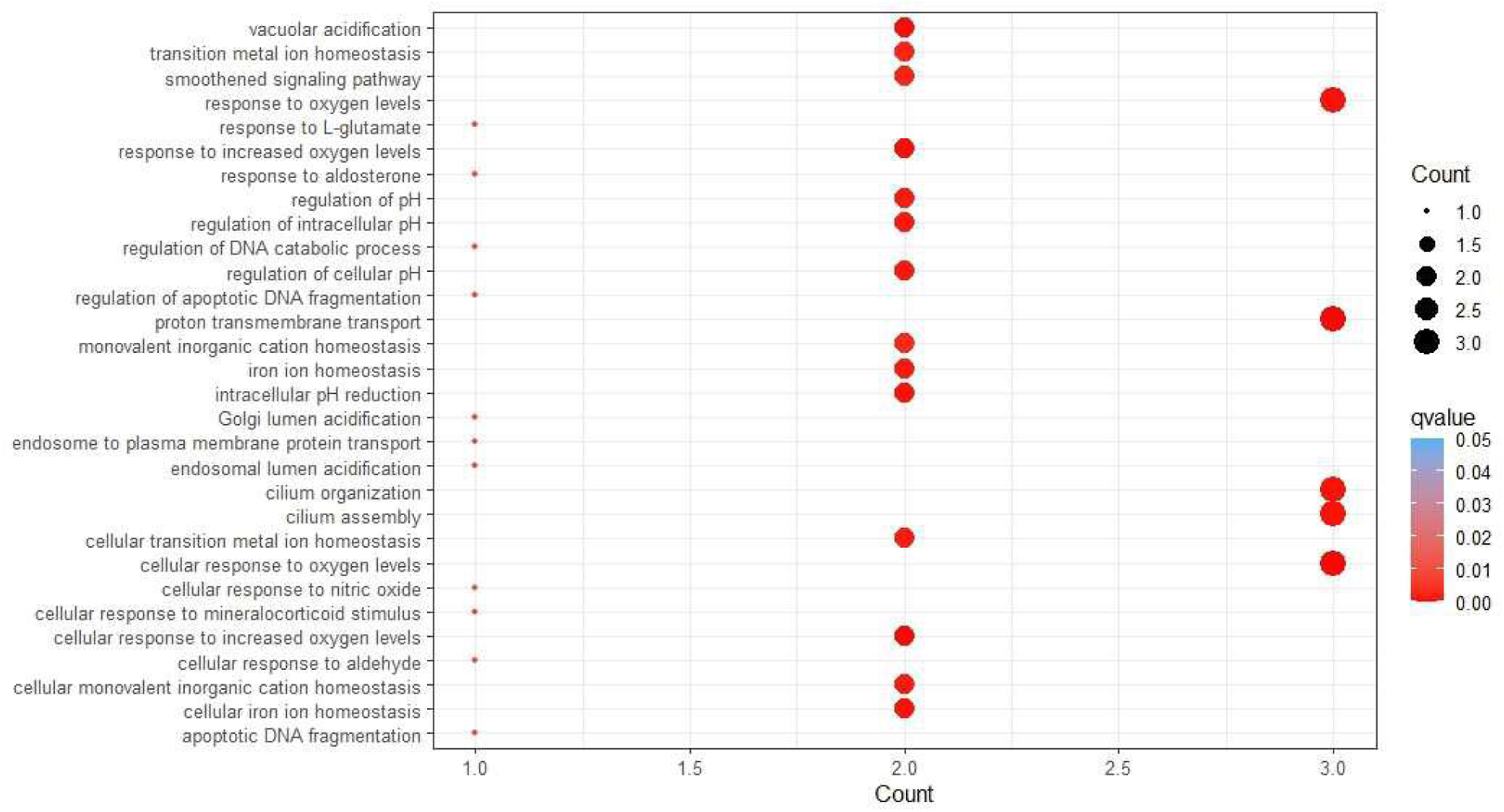
Dot plot representing top 30 pathways in the brain influenced by downregulated proteins interacting with SARS-CoV-2 NSP6 interacting proteins. Dot size represents the gene ratio and dot color depicts the q-value as in the heat map. X-axis shows the count of genes per GO term and Y-axis indicates the GO terms. The color gradient indicates the *q*-value using the Benjamini-Hochberg method.

In the case of the heart, the influenced pathways by upregulated genes were viral processes, cellular cation homeostasis, viral life cycle, protein targeting, myeloid leukocyte activation, macroautophagy, etc. (Figure 9a and Supporting Information Data 5).

**Figure 9a.**
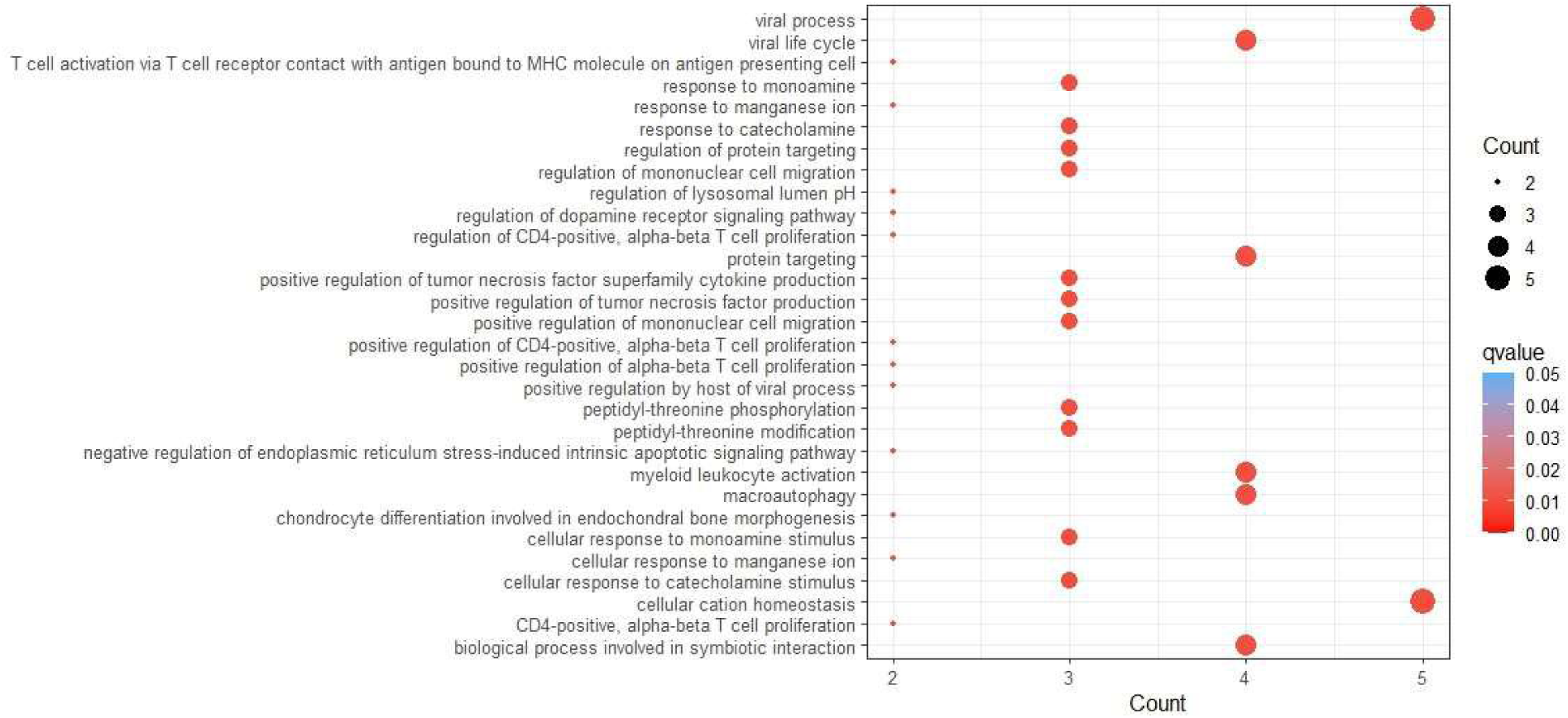
Dot plot representing top 30 pathways in the heart influenced by upregulated proteins interacting with SARS-CoV-2 NSP6 interacting proteins. Dot size represents the gene ratio and dot color depicts the q-value as in the heat map. X-axis shows the count of genes per GO term and Y-axis indicates the GO terms. The color gradient indicates the q-value using the Benjamini-Hochberg method.

The down regulated genes were observed to act on cilium assembly and organisation, cellular response to oxygen levels, protein targeting, negative regulation of organelle organisation and cell cycle process, etc. (Figure 9b and Supporting Information Data 5). None of the pathways had a significant q-value.

**Figure 9b.**
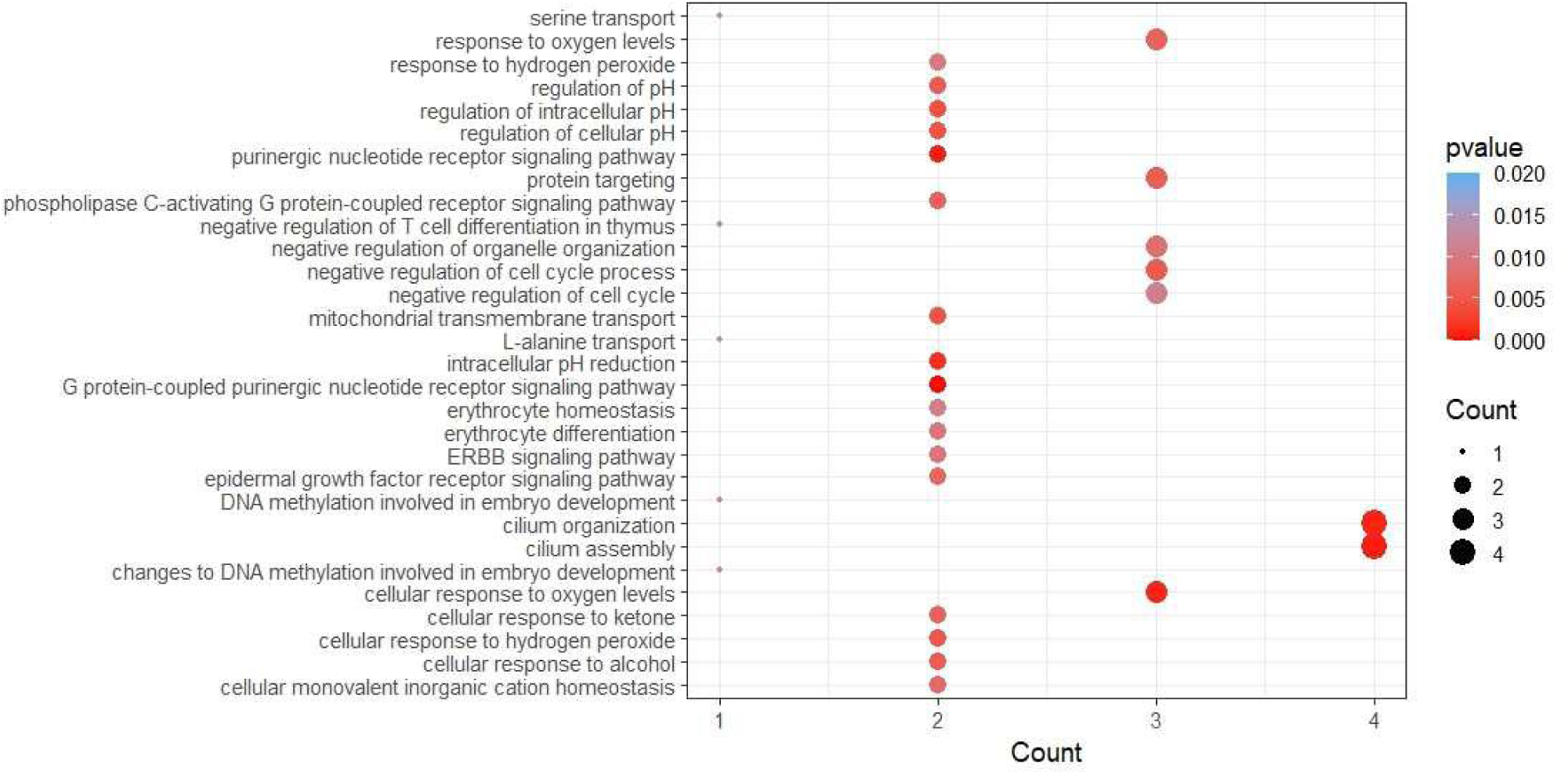
Dot plot representing top 30 pathways in the heart influenced by downregulated proteins interacting with SARS-CoV-2 NSP6 interacting proteins. Dot size represent the gene ratio and dot color depicts the p-value as in the heat map. X-axis shows the count of genes per GO term and Y-axis indicates the GO terms. The color gradient indicates the p-value.

Within the lung, the upregulated genes had been noticed to influence some pathways like viral life cycle and viral entry into host cell, tumor necrosis factor production, stress-activated protein kinase signalling cascade, stress activated MAPK cascade, response to interferon gamma, regulation of regulated secretory pathway, exocytosis regulation, neurotransmitter transport and secretion, signal release from synapse, myeloid leukocyte activation, regulation of biological process involved in symbiotic interaction, modulation by symbiont of entry into host, movement in host, etc. (Figure 10a and Supporting Information Data 5).

**Figure 10a.**
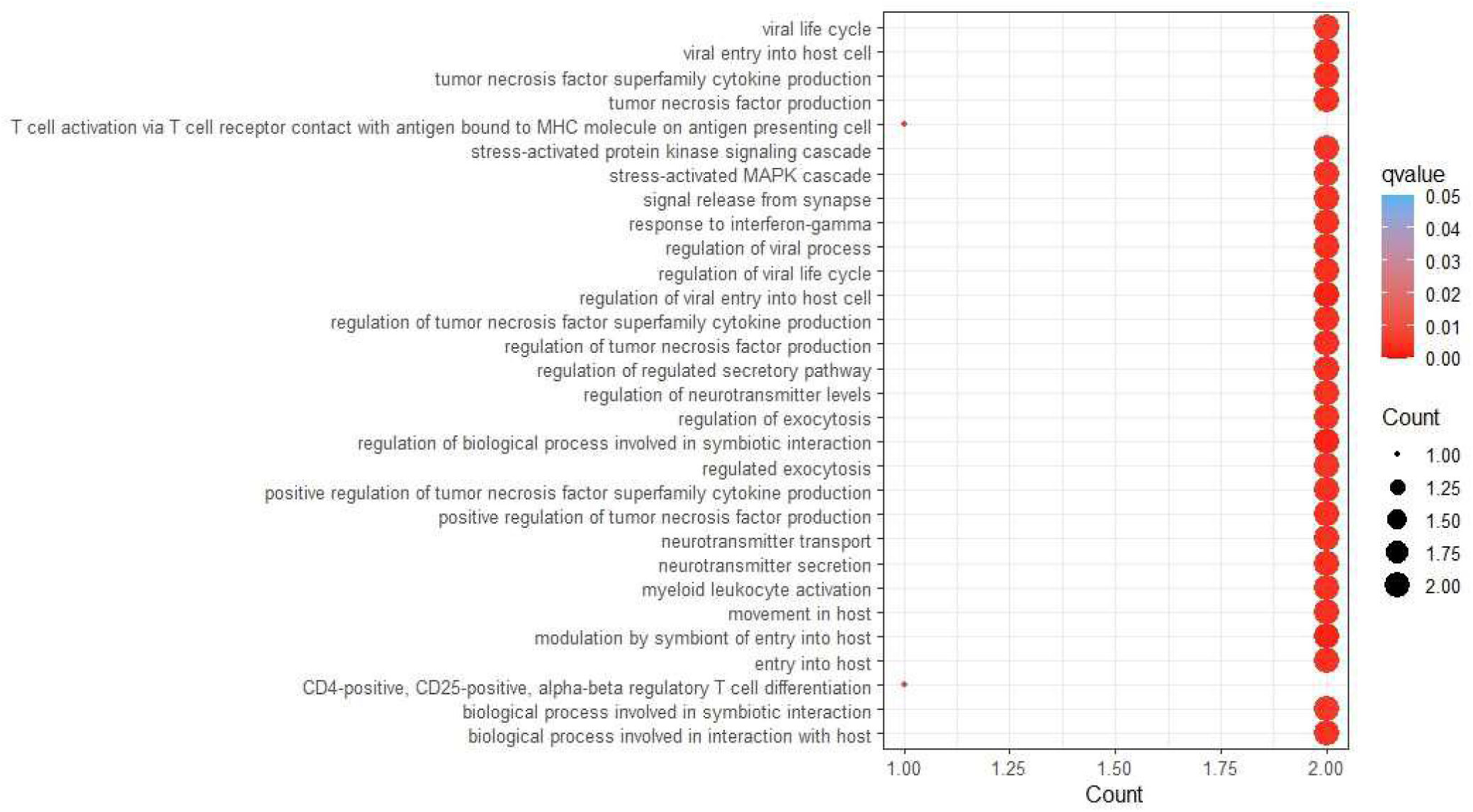
Dot plot representing top 30 pathways in the lung influenced by upregulated proteins interacting with SARS-CoV-2 NSP6 interacting proteins. Dot size represents the gene ratio and dot color depicts the q-value as in the heat map. X-axis shows the count of genes per GO term and Y-axis indicates the GO terms. The color gradient indicates the q-value using the Benjamini-Hochberg method.

In the case of pathways influenced by genes down regulated in the lung, we found response to oxygen levels, positive regulation of protein localization, positive regulation of protein transport, synapse organisation, vacuolar acidification, T cell proliferation, regulation of synaptic plasticity, regulation of pH, regulation of macroautophagy, regulation of mononuclear cell and lymphocyte proliferation, regulation of endocytosis, monovalent inorganic cation homeostasis, intracellular pH reduction, etc. (Figure 10b and Supporting Information Data 5).

**Figure 10b.**
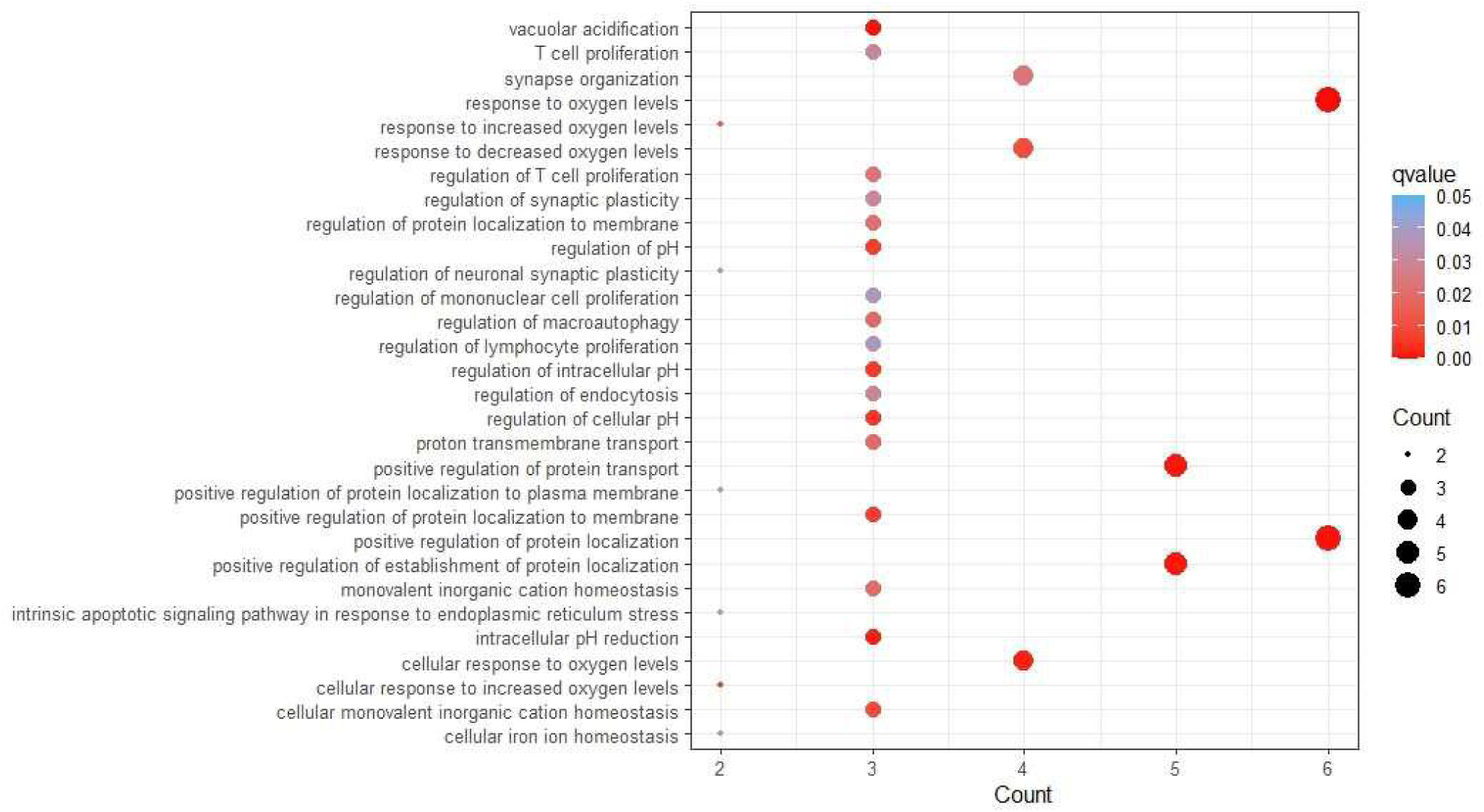
Dot plot representing top 30 pathways in the lung influenced by downregulated proteins interacting with SARS-CoV-2 NSP6 interacting proteins. Dot size represents the gene ratio and dot color depicts the q-value as in the heat map. X-axis shows the count of genes per GO term and Y-axis indicates the GO terms. The color gradient indicates the q-value using the Benjamini-Hochberg method.

### 3.7. SARS-CoV-2 influenced miRNA population is different in the brain and lung

Protein regulation by miRNAs could affect protein abundance in cells and influence pathways where these proteins are involved. After finding out the different pathways in the brain, heart, and lung we looked for miRNAs expressed in these tissues that can target interacting partners of SARS-CoV-2 NSP6 interacting proteins regulated by SARS-CoV-2 infection using miRNet. We found 82 miRNAs in the brain and 43 miRNAs in the lung. Among these 16 were common in the brain and lung. 66 miRNAs were unique for brain whereas for lung it was 27 (Figure 11 and Supporting Information Data 6). In miRNet, miRNAs expressed in the heart were not available.

**Figure 11.**
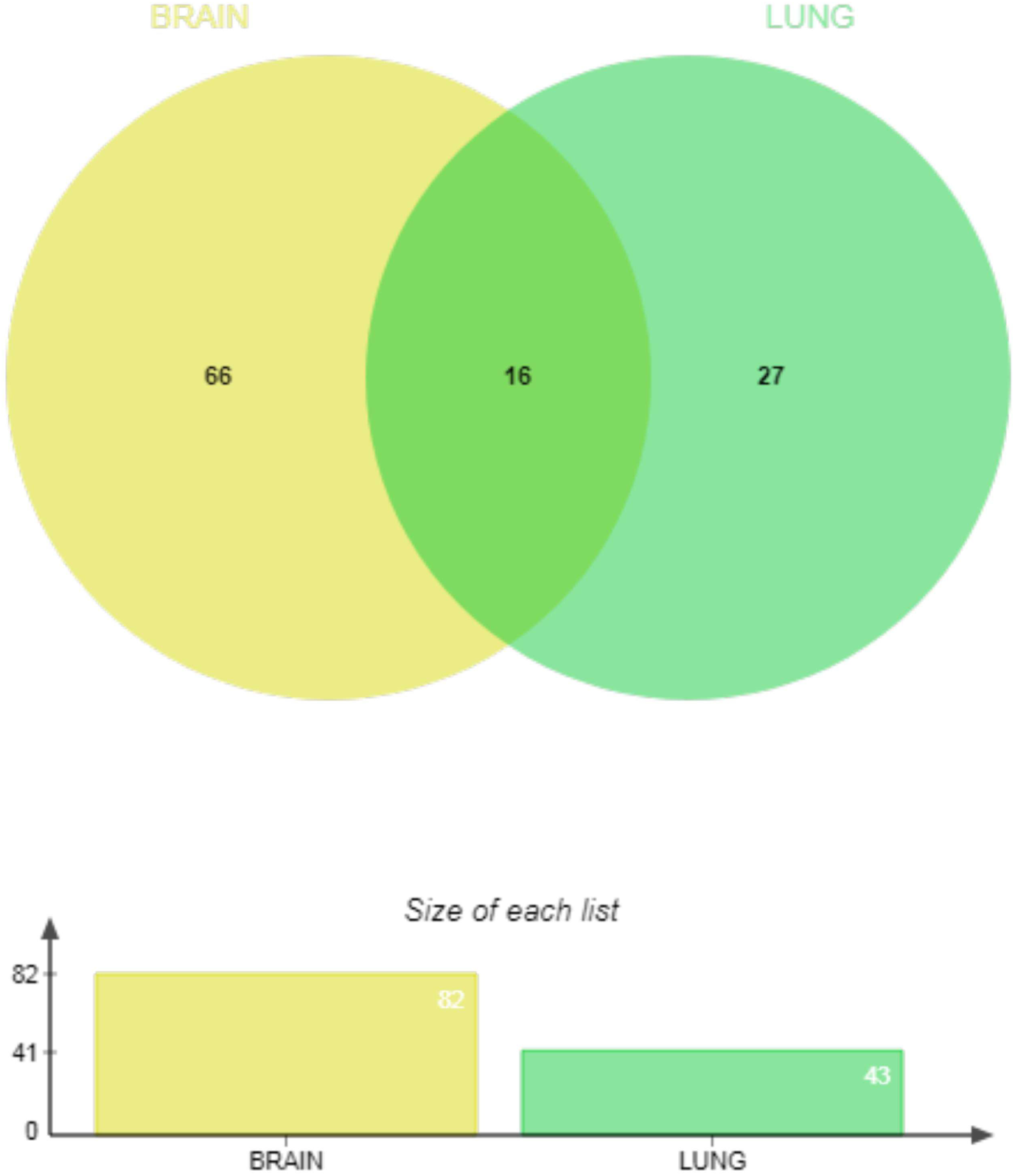
Common miRNAs in the brain and lung. Proteins influenced by SARS-CoV-2 infection that interact with SARS-CoV-2 NSP6 in the brain and lung were taken to find out miRNAs that can target these protein mRNAs. Among the total of 82 miRNAs in the brain and the total of 43 miRNAs in the lung, 16 miRNAs were common whereas 66 and 27 miRNAs were unique in the brain and lung respectively.

#### 3.8.1. The miRNA-mRNA interaction network involving the brain and lung expressed miRNAs targeting SARS-CoV-2 influenced interacting partners of SARS-CoV-2 NSP6 interacting proteins and their corresponding pathways

From gene-microRNA interaction network analysis of the brain, the gene-microRNA network consisted of 113 nodes, (82 miRNAs, 18 genes, and 13 influencing transcription factors (TF)), and 297 edges. In the miRNA interaction network, we identified five miRNAs with comparatively high degree (≥10) and betweenness values (hsa-mir-124-3p with degree 14 and betweenness 484.1903, hsa-mir-1-3p with degree 13 and betweenness 431.2286, hsa-mir-16-5p with degree 12 and betweenness 379.2163, hsa-let-7b-5p with degree 11 and betweenness 239.3807, and hsa-mir-155-5p with degree 10 and betweenness 243.6051) as depicted in figure 12. Pathways influenced by these miRNAs are given in supporting information data 7 that were obtained from GO-BP and KEGG pathways. These pathways included transcription related pathways, pathways regulating cell proliferation, pathways regulating nucleotide metabolism, apoptotic signalling pathways, regulation of myeloid cell and leukocyte differentiation, etc., in the GO-BP list. KEGG list consisted of pathways (p≤ 0.05) in viral infection (Epstein-Barr virus, HTLV-I, Hepatitis C, and Herpes simplex) various cancer pathways (chronic myeloid leukemia, acute myeloid leukemia, colorectal cancer, and prostate cancer), different signalling pathways (MAPK signalling, Wnt signalling, p53 signalling, and ErbB signalling), and pathways in bacterial infection (pertussis and tuberculosis) (Supporting Information Data 8).

**Figure 12.**
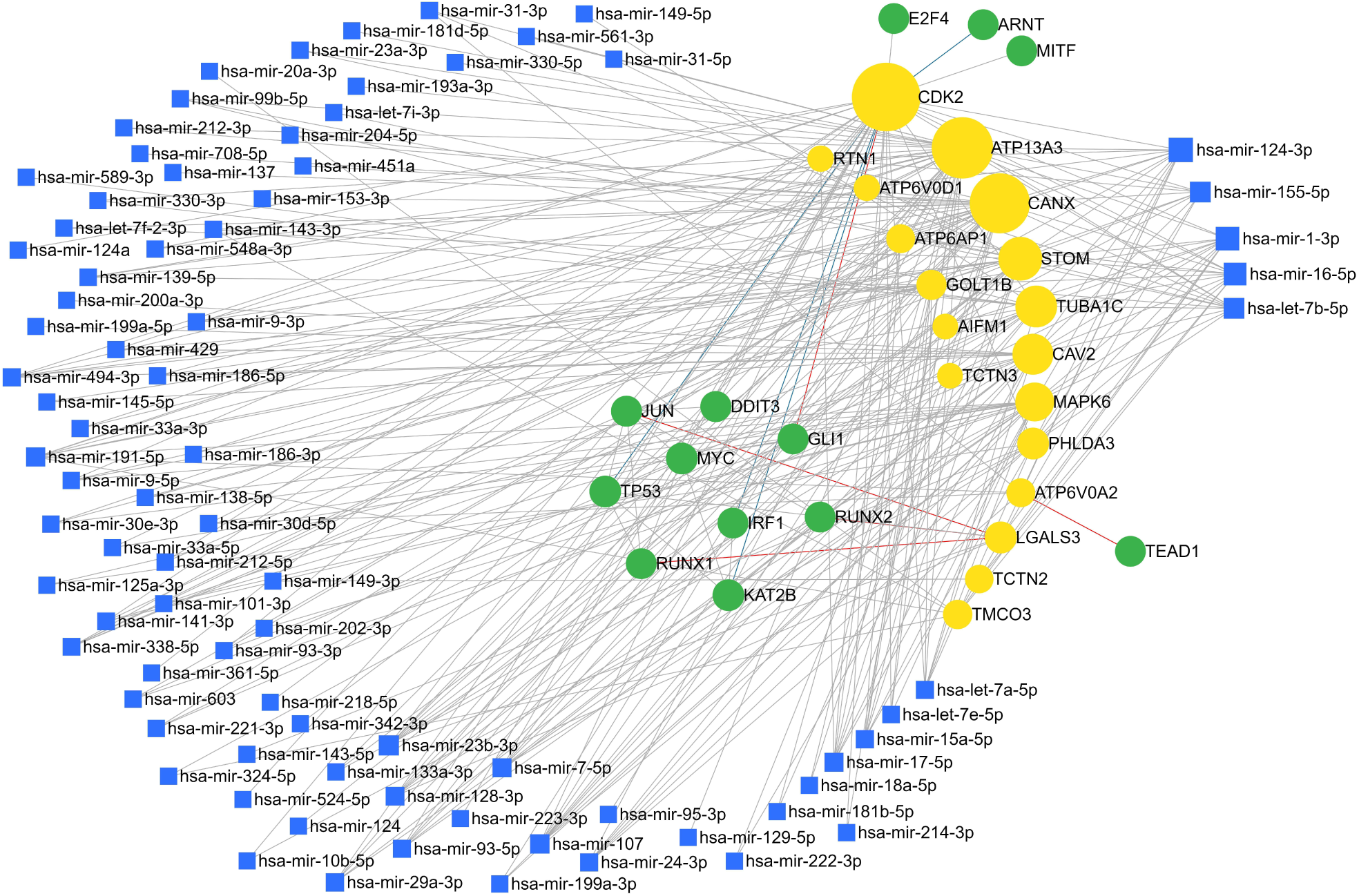
miRNA interaction network with interacting proteins of SARS-CoV-2 NSP6 interacting proteins in the brain. Yellow colored nodes denote proteins and green colored nodes are transcription factors influencing proteins and miRNAs in the network. Blue squares represent miRNA nodes. The increased size of node represents a higher degree.

In the lung tissue, 108 nodes consisting 43 miRNAs, 22 genes, and 43 TFs exhibited 337 edges. In the lung specific miRNA interaction network, we identified two miRNAs with comparatively high degree (≥10) and betweenness values (hsa-mir-1-3p with degree 14 and betweenness 403.7072 and hsa-mir-34a-5p with degree 10 and betweenness 569.9464) (Figure 13 and Supporting Information Data 9). This indicated the significance of these miRNAs as hubs (for their high degree) and their important role in the interaction network (for their high betweenness). Influenced pathways by these miRNAs are given in supporting information data 10 obtained from GO-BP and KEGG. GO-BP pathway list consisted of regulation of DNA-dependent transcription, regulation of RNA metabolic processes, positive regulation of transcription from RNA polymerase II promoter, regulation of cell differentiation, cell proliferation, organ development, embryo development, nervous system development, etc. Pathways with p-value≤ 0.05 in KEGG pathway list were HTLV-1 infection, cancer pathways (prostate, renal cell, pancreas, colorectal, and chronic myeloid leukemia), osteoclast differentiation, prion diseases, signalling pathways (TGF-beta, ErbB, Notch, Wnt, and MAPK), tuberculosis, addiction (cocaine, amphetamine, and alcoholism), Leishmaniasis, mineral absorption, and adherens junction.

**Figure 13.**
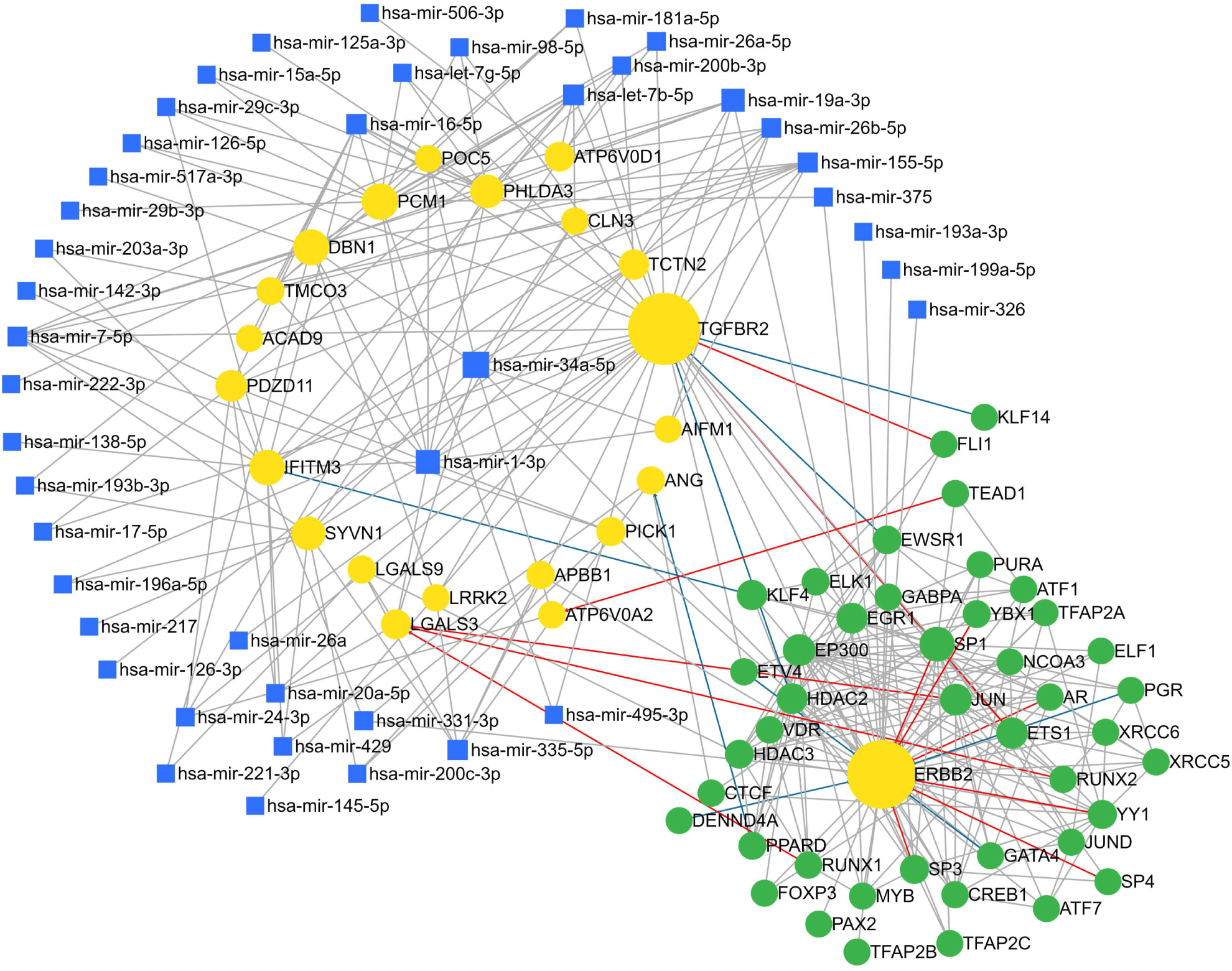
miRNA interaction network with interacting proteins of SARS-CoV-2 NSP6 interacting proteins in the lung. Yellow colored nodes denote proteins and green colored nodes are transcription factors influencing proteins and miRNAs in the network. Blue squares represent miRNA nodes. The Increased size of node represents a higher degree.

#### 3.8.2. Possible pathways in heart influenced by SARS-CoV-2 regulated circulating miRNAs in the heart

In miRNet heart specific miRNA database is absent. To construct probable miRNA-RNA interaction network of heart that is influenced by SARS-CoV-2 NSP6, we took miRNAs and genes that satisfied two conditions – i) SARS-CoV-2 regulated circulating miRNAs^56,58^ and (ii) among this list, miRNAs targetting SARS-CoV-2 regulated proteins in the heart interacting with SARS-CoV-2 NSP6 interacting proteins. Resulted miRNA-RNA interaction network is shown in figure 14 and supporting information data 11.

**Figure 14.**
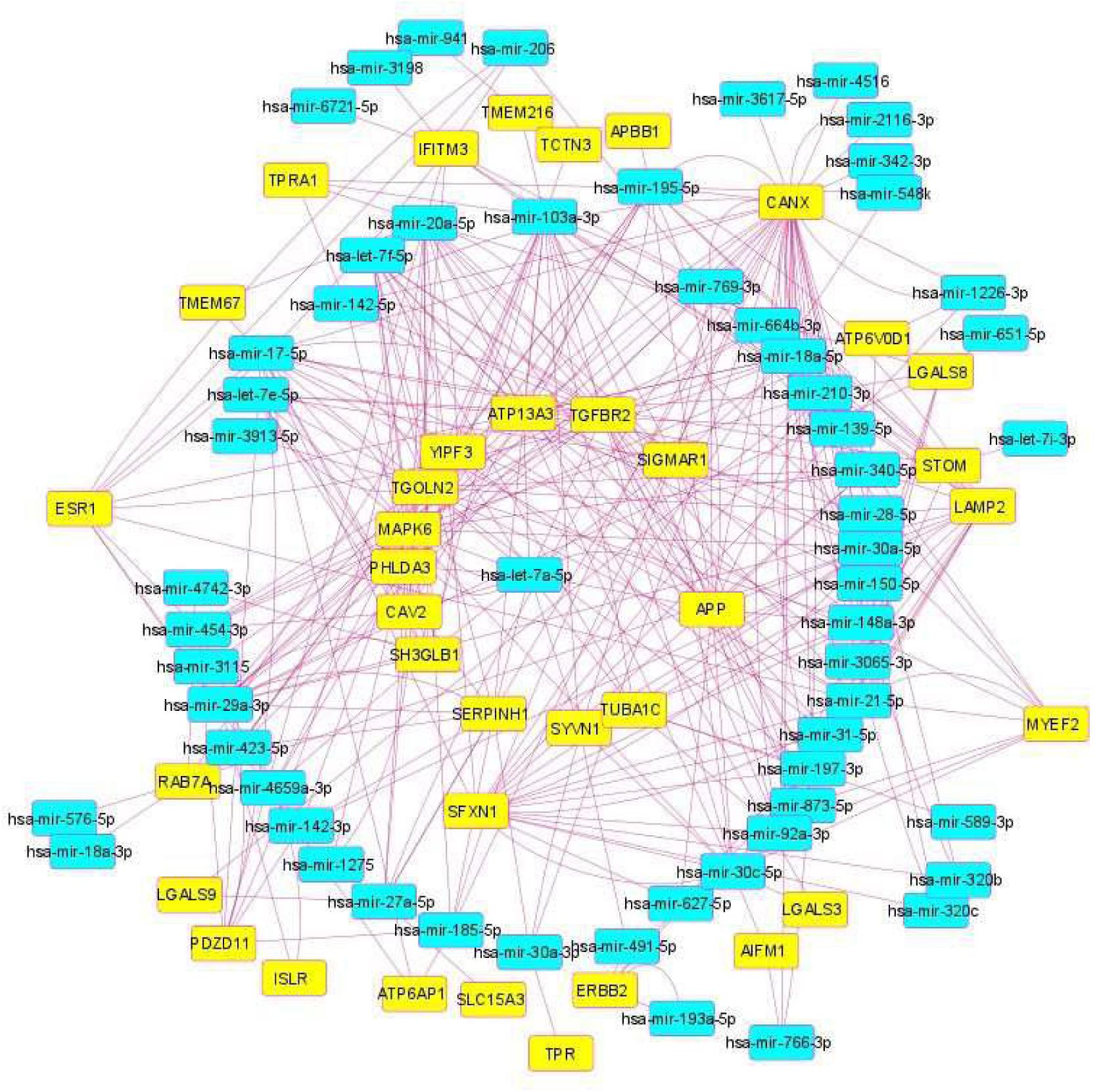
miRNA interaction network with interacting proteins of SARS-CoV-2 NSP6 interacting proteins in the heart. Yellow colored nodes denote proteins and blue colored nodes are miRNAs in the network.

### 3.9. Regulated SARS-CoV-2 influenced circulatory miRNAs were found to be targeting tissue-specific interacting proteins of SARS-CoV-2 NSP6 interacting proteins

As SARS-CoV-2 is reported to be influencing miRNAs^56,58^, we wanted to study the regulated circulatory miRNAs and miRNAs that target SARS-CoV-2-regulated proteins interacting with SARS-CoV-2 NSP6 binding proteins in the brain and lung. Common miRNAs in the reported list of SARS-CoV-2-influenced circulating miRNAs^56,58^ and tissue-specific miRNAs influencing interacting proteins of SARS-CoV-2 NSP6 binding proteins were probed to find out regulated miRNAs and their respective target proteins. Among the common miRNAs in the lung we observed three miRNAs (hsa-mir-17-5p, hsa-mir-20a-5p, and hsa-mir-142-3p) whereas, in the brain it was ten miRNAs (hsa-let-7a-5p, hsa-let-7e-5p, hsa-mir-17-5p, hsa-mir-18a-5p, hsa-mir-29a-3p, hsa-mir-31-5p, hsa-mir-139-5p, hsa-mir-342-3p, hsa-mir-589-3p, and hsa-let-7i-3p) (Supporting Information Data 12). With the help of miRNET, we found out genes targeted by these miRNAs in the brain and lung. In the lung 1616 genes were targeted by the three observed miRNAs (Figure 15 and supporting information data 13) and 2782 genes were observed to be targeted by the 10 miRNAs in the brain (Figure 16) (Supporting Information Data 14). We wanted to see the common proteins that were targeted by the regulated miRNAs (both in the brain and lung). We found 3 genes common in the brain and 2 genes common in the lung. Checking for the pathways that were involving these genes, we were able to see pathways like induction of tolerance to self antigen (B cell, T cell, NK T cell), protein amino acid phosphorylation, protein metabolic process, phosphorous metabolism, regulation of mitosis, response to cholesterol, myeloid dendritic cell differentiation, etc., (Figure 17) (Supporting Information Data 15).

**Figure 15.**
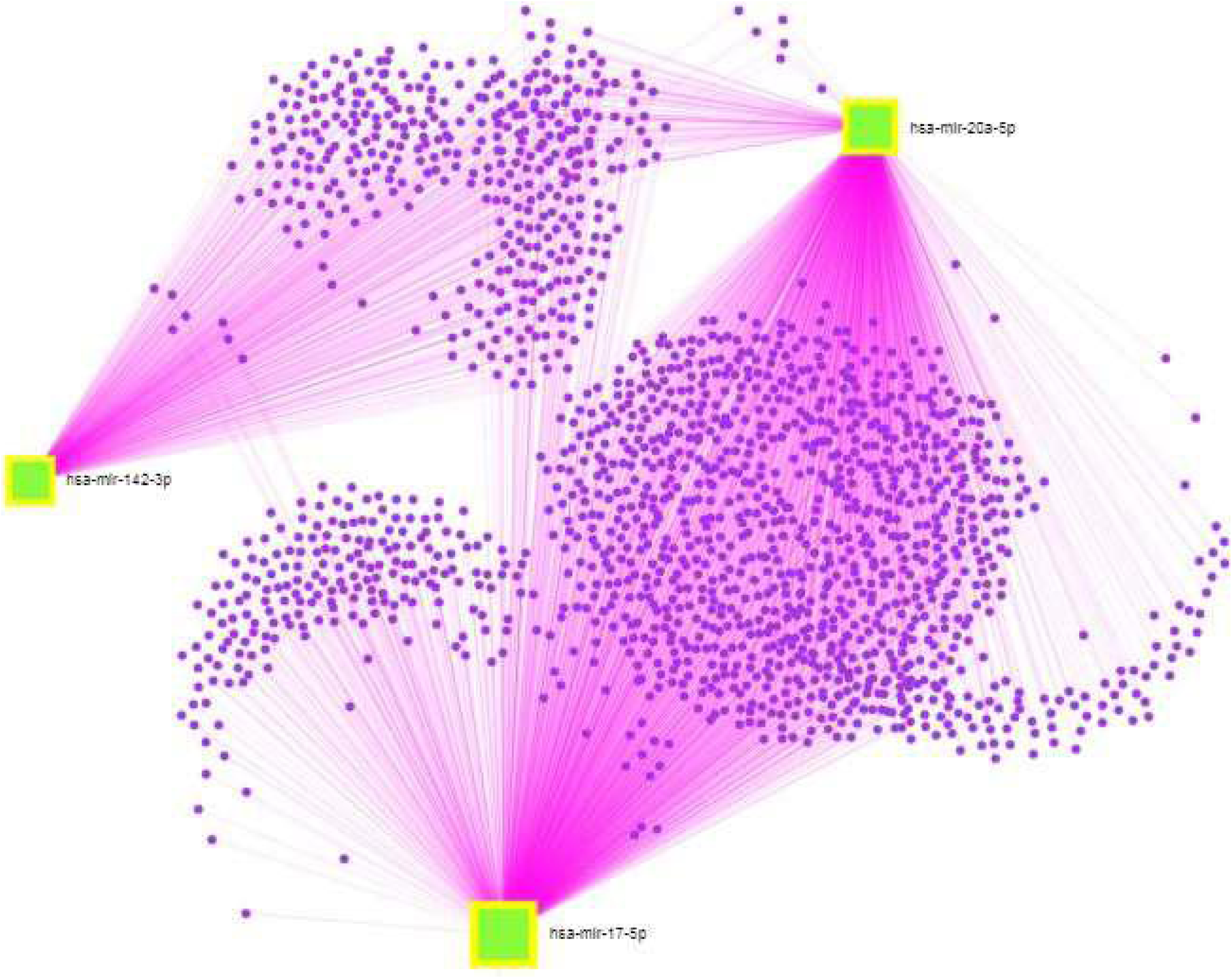
Common among SARS-CoV-2 influenced circulating miRNAs and lung miRNAs. Square nodes denote miRNAs and circular nodes represent targeted mRNAs. Three miRNAs were targeting 1616 mRNAs.

**Figure 16.**
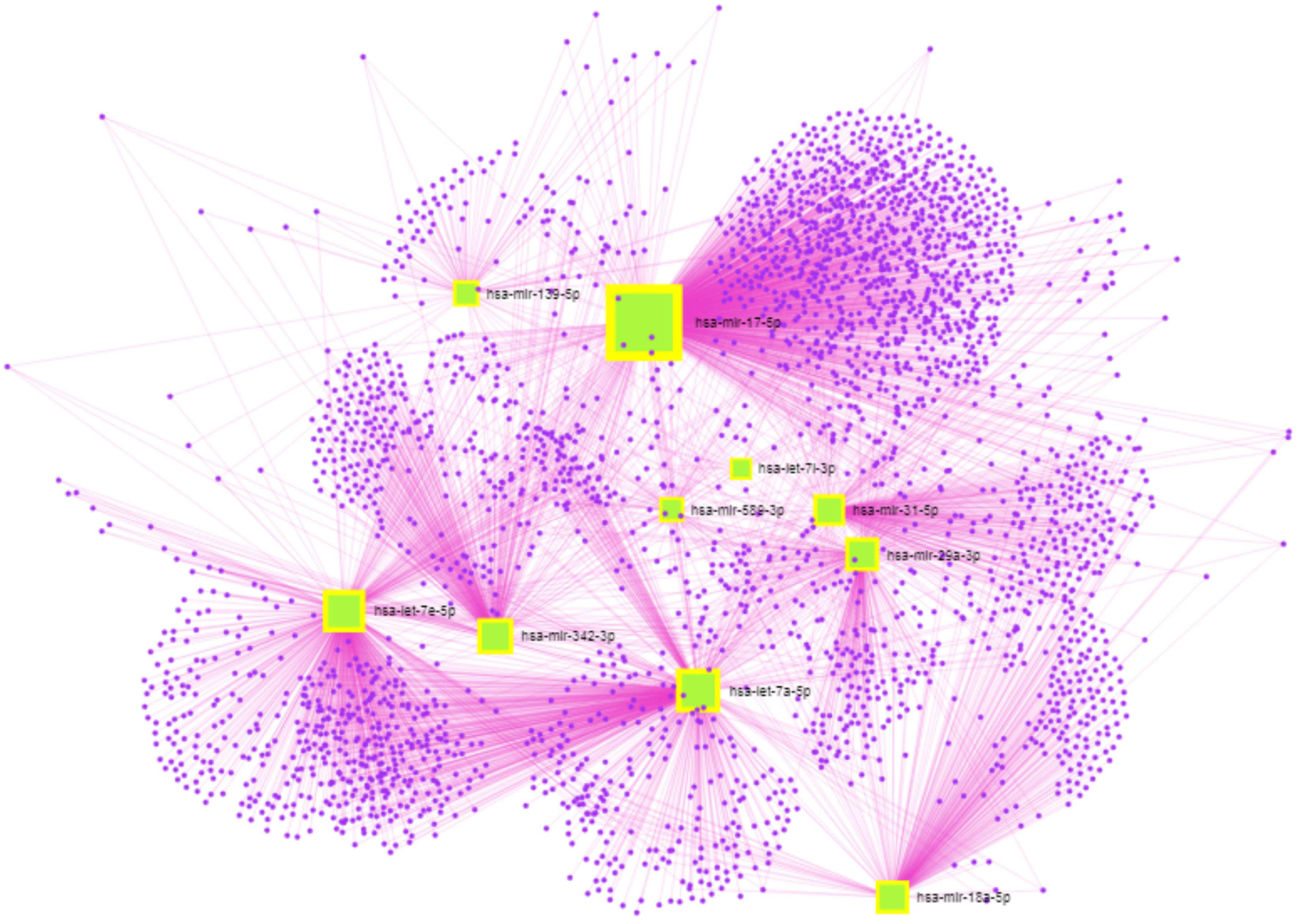
Common among SARS-CoV-2 influenced circulating miRNAs and brain miRNAs. Square nodes denote miRNAs and circular nodes represent targetted mRNAs. 10 miRNAs were targeting 2782 genes.

**Figure 17.**
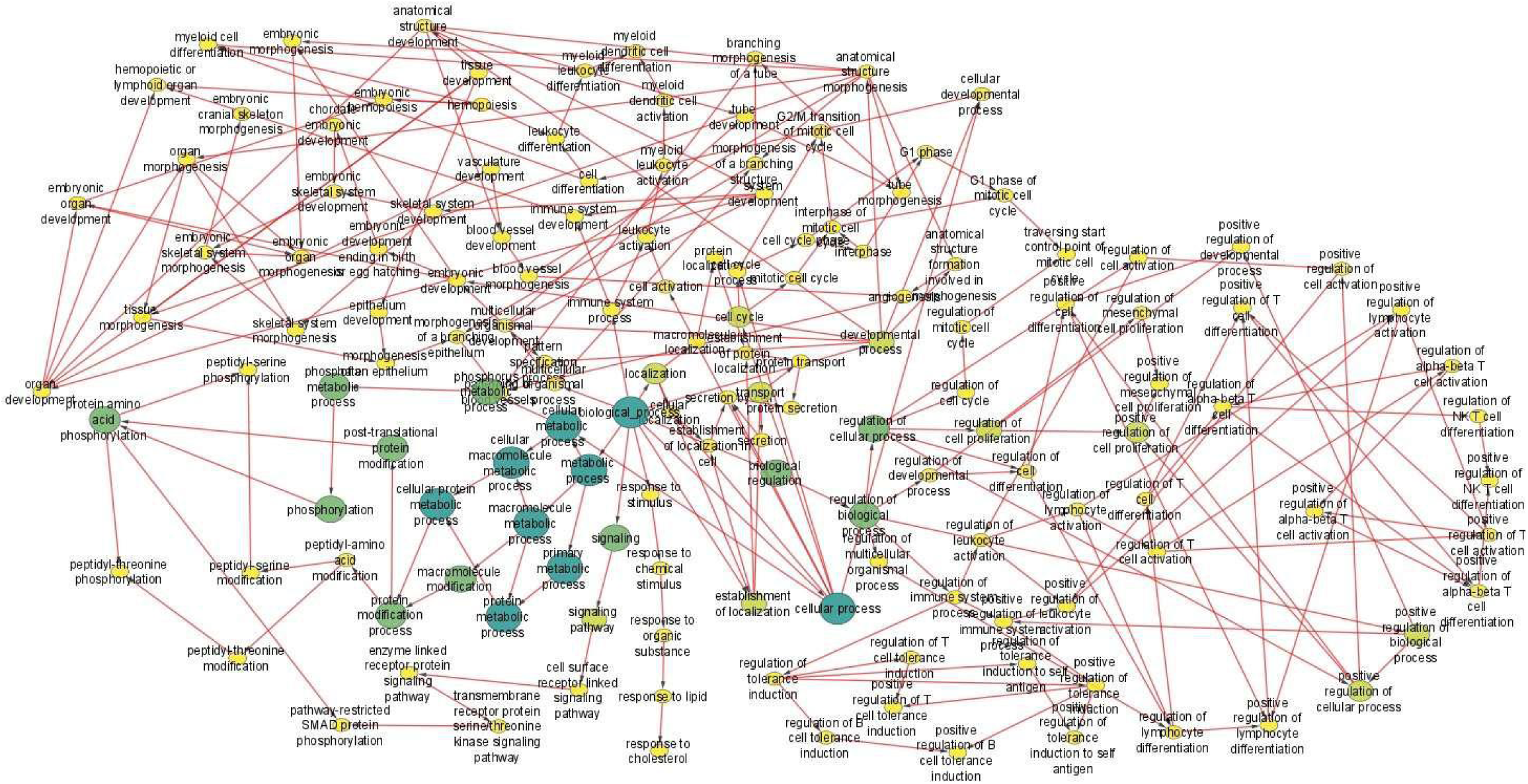
Pathways involving common 5 genes observed from the brain and lung. Circular nodes represent pathways associated with 5 genes. Continuous color mapping from green to yellow represents maximum to minimum significance.

## 4. Conclusions and Discussion

In various analyses, tissue-specific complications due to SARS-CoV-2 infection have been delineated, but in the case of individual influence of SARS-CoV-2 accessory proteins they are yet to be delineated in detail. Here we have taken SARS-CoV-2 NSP6 protein to find its tissue-specific host response. Proteins influenced after SARS-CoV-2 infection in the brain, heart, and lung that are interacting with SARS-CoV-2 NSP6 interacting proteins were taken for finding pathways influenced by these proteins as well as the hub genes that could be important players in host tissue-specific pathogenesis. Clue GO analyses provided a broad picture of probable pathways and biological processes undergoing perturbations related to these hub genes in the brain, heart, and lung of COVID infected individuals. In addition, reported circulatory miRNAs regulated after SARS-CoV-2 infection were also taken for miRNAs expressed in the brain and lung to find out miRNA-mRNA network and miRNA influenced pathways in these organs. To find out possible miRNA influenced pathways in the heart, SARS-CoV-2 regulated genes targeted by reported circulatory miRNAs regulated after SARS-CoV-2 infection were used.

Among the common proteins in the brain, heart, and lung, that were influenced by SARS-CoV-2, AIFM1 was reported to be involved in apoptosis,^61^ and in COVID-19, anti-neuronal autoantibody against this protein was found to be formed^62^. This protein was shown to be involved in central respiratory drive and reflex responses in impaired lung function^62^. ATP6V0D1 was observed to be upregulated in A549 cells transfected with SARS-CoV-2 S plasmid and intracellular/lysosomal pH was observed to be decreased.^63^ TCTN2, another regulated protein was shown to be associated with osteoarthritis.^64^ PHLDA3 knock down was observed to be negatively affecting colony forming efficiency of MM468 cells expressing myr-Akt and high frequency of chromosome loss at PHLDA3 locus was also reported in large cell neuroendocrine carcinoma (LCNEC).^65^

Our study of hub genes influenced pathways in the brain included pathways observed in cancer like small cell lung cancer (CDK2, CDKN1B, CCNE1, CCND1), and prostate cancer (CDK2, CDKN1B, CCNE1, CCND1) and these genes identified could be indirectly linked to genes regulating dysregulation of cell cycle, it’s repair, and checkpoints. With the hub genes regulated by SARS-CoV-2 infection affecting these pathways could have an effect in these cancers. It is reported that NSP6 and ORF7a through transforming growth factor ß-activated kinase 1 (TAK1) activate NF-ĸB which in turn promotes proinflammatory cytokine levels.^66^ In lung cancer CCND1 overexpression by NF-ĸB translocation was observed and was shown to be promoting cancer through PI3K/AKT signaling pathway.^67,68^ Protein-protein interactome with NSP6 interacting partners in the brain exhibited CCND1, CDK2, and CCNA1 as crucial players mainly altering cell cycle processes. The majority of the pathway terms were G1/S transition and the next highest group consisted of iron uptake and transport terms. Activation of PTK6 and regulation of cell cycle associated pathways, and p53 signalling pathways were also found in the analysis. This instigates for further exploration and elucidation of impact of coronavirus NSP6 on cancer. For the lung hub genes influenced pathways, ATP6V0A1, ATP6V0A2, ATP6V0D1, ATP6V1D, ATP6V1B1, and ATP6V1E1 were involved in intraphagosomal pH lowering by the V-ATPase. SARS-CoV-2 was shown to be increasing the expression of V-ATPase.^63^ Pathway involving other hub genes TGFB1, TGFBR2, and LGALS3 was dendritic cell differentiation. Further analyses of pathways involving heart hub genes RAB7A, ATPV0D1, and LAMP2 were associated with phagosome related processes.

Another hub gene TGFB1 showed association with rheumatoid arthritis that is also one of the reported disease in postacute sequelae of COVID-19.^69^ Hub genes in the brain CDK2, CCNB1, CCNE1, CCND1, and CDKN1B are drug targets for both approved and non-approved drugs with some of them having more than one target protein. Some approved drugs were targeting multiple proteins in our analyses. Acetaminophen targets CDK2 and CCND1, palbociclib targets CCND1 and CCNE1, raltitrexed targets CDK2 and CDKN1B, lapatinib targets CCND1 and CDKN1B, and doxorubicin hydrochloride, lapatinib, and methotrexate target both CCND1 and CDKN1B. Considering the hub genes in heart, TGFB1, CANX, APP, and LGALS3 were the drug targets and one of the drug inositol targets both TGFB1 and APP. In the lung, among the drug targets TGFB1, LGALS3, TGFBR2, and EGF, irinotecan hydrochloride targets both TGFB1 and TGFBR2. Drugs targeting hub genes were anti-cancer drugs like palbociclib, lapatinib, irinotecan, etc., anti-rheumatic like methotrexate, anti-pyretic like acetaminophen, etc.

Deleterious effect of NSP6 in the heart or cardiomyocytes is reported. Expression of NSP6 in the *Drosophila* heart caused cardiac actin filaments getting disorganized and significantly reducing fiber density affecting its function with reduced diastolic diameter of the heart tube.^70^ SARS-CoV-2 NSP6, NSP8 and M overexpression caused transcriptome and proteome change in human pluripotent stem cell (hPSC) derived cardiomyocytes as well as undergo apoptosis.^37^

We found galectin-3 (LGALS3), one of the binding partners of SARS-CoV-2 NSP6 interacting proteins regulated in COVID-19 infection. It was observed to be upregulated in the heart after SARS-CoV-2 infection. Myocardial galectin-3 expression was reported to be significantly correlated with inflammatory cell count on endomyocardial biopsy in inflammatory cardiomyopathy.^71^ Galectins are reported to be tumor progression modulators.^72,73^ Among the 10 updated “hallmarks of cancer”^74^ galectin-3 was reported to be influencing all of them directly or indirectly in a number of tumors.^73^ In the context of “sustaining proliferative signalling” galectin-3 was reported to be binding to activated K-RAS and influencing tumor cell proliferation and progression.^75,73^ Mutant p53 was shown to loose transcriptional repression of galectin-3^76^ as well as loss of p53 cooperating protein HIPK2 (homeodomain-interacting protein kinase 2) showed increased galectin-3 expression in well differentiated thyroid carcinomas.^73,77^ Knock down of galectin-3 arrested prostate cancer cells at G1 phase.^78^ These aspects show the influence of galectin-3 in another hallmark of cancer, ie., “evading growth suppressors”. Reports on galectin-3 promoting CTL dysfunction,^73,79^ and limiting NK cell attack on tumor^73,80^ show it’s influence on “immune destruction avoidance”. Galectin-3 knockdown and thereby decreased hTERT expression leading to induction of cellular senescence in gastric cancer cells as well as galectin-3 interaction with hTERT through it’s N-terminal (1 – 63 aa)^81^ shows it’s influence on another hallmark of cancer, “enabling replicative telomerase”. With respect to “tumor promoting inflammation”, galectin-3 was reported to be inducing the release of pro-inflammatory cytokines that promote metastasis.^82^ Increased amount of circulating galectin-3 was reported in metastatic colorectal cancer (31.6 fold) in comparison to non-metastatic patients (11.3 fold) and healthy individuals.^78^ Also it was reported to be increased in breast cancer patients (11.3 fold) with respect to healthy individuals.^83^ In addition, increased level of galactin-3 in comparison to healthy individuals was also reported in gastrointestinal, lung, and ovarian cancers and non-Hodgkin’s lymphoma with metastatic cases showing even higher levels.^84^

Increased level of circulating galectin-3 was also observed in melanoma patients and patients with greater than 10ng/mL level had the worst outcome in comparison to less than 8ng/mL group and 8-10ng/mL group.^85^ Mechanistically galectin-3 was reported to be stimulating secretion of tumor promoting interleukin (IL)-6, granulocyte colony-stimulating factor (G-CSF), sICAM-1, and granulocyte macrophage colony-stimulating (GM-CSF) from blood vascular endothelial cells both *in vitro* and in *in vivo* mouse study leading to upregulation of metastasis associated adhesion molecules integrin α_V_β_1_, vascular cell adhesion molecule-1, and E-selectin.^82^ This also points to the involvement of galectin-3 in the “activation of metastasis and invasion”. Various roles of galectin-3 reported to be influencing activation of metastasis and invasion are (i) enhancing cell detachment through secreted galectin-3 as it weakens the attachment of tumor cell adhesion molecules to extracellular matrix proteins like laminin and fibronectin,^86^ (ii) enhance binding of breast cancer cells expressing cancer-associated Thomsen-Friedenreich antigen (T antigen) to galectin-3 surface expressing endothelial cells,^87,88^ (iii) galectin-3 binding to core1 antigen of cancer cells causes homotypic aggregation of low alpha-N-acetylgalactosaminide alpha-2,6-sialyltransferase 2 (ST6GalNAc2) expressing cancer cells and have low anoikis-mediated apoptosis and heterotypic adhesion to endothelium promoting metastasis.^89^ In the induction of angiogenesis, galectin-3 was shown to be a pro-angiogenic molecule and a mediator of vascular endothelial growth factor (VEGF) and basic fibroblast growth factor (bFGF) mediated angiogenic response by modulating αvβ3 integrin activity.^90^ Galectin-3 was also reported to be acting as molecular regulator of Jagged-1 (JAG1)/Notch signalling pathway of angiogenesis.^91^ In the context of contribution of galectin-3 in “acquiring genomic stability”, it was reported to be interacting with DNA damage repair (DDR) protein like BARD1,^92,93^ which is the main partner of breast and ovarian cancer susceptibility gene product BRCA1, where both the proteins are involved in homologous recombination (HR) based DDR. Galectin-3 was also reported to be interacting with four DDR-related proteins namely PARP1, HSP90AB1, CDC5L, and PRPF19 and its knock down increased DNA damage resistance in HeLa cells.^93^ Developing resistance to cell death, another hall mark of cancer is important for the survival of cancer cells. Galectin-3 was observed to be influencing cell death depending upon the target protein it binds.^94^ In human B and T cell lines, endogenous galectin-3 binding to CD95 (APO-1/Fas) acts as pro-apoptic, whereas C2-ceramide-mediated apoptosis was inhibited.^95^ In bladder cell carcinoma cell line J82, galectin-3 by increasing Akt activity, inhibited TNF-related apoptosis-inducing ligand (TRAIL) induced apoptosis.^96^ Human breast carcinoma cell line BT549 cells became susceptible to TRAIL induced apoptosis after galectin-3 cDNA expression.^97^ TRAIL induced apoptosis-resistant metastatic colon cancer cell line LS-LIM6 was found to be having increased cell surface expression of galectin-3 and its silencing or blocking using inhibitors and glycosylation conferred sensitivity again to cell death showing anti-apoptotic function of galectin-3.^98^ Cancer cells need to be changing their energy metabolism (metabolic reprogramming) unlike normal cells, which would help them provide protein, lipid and nucleic acids necessary for accelerated division rate that is another hallmark of cancer. Regulation of glycogen synthase kinase-3β activity by galectin-3,^99^ increased glutamate-driven tricarboxylic acid cycle by collagen deposition contributed by increased galecting-3 secretion^100^ could be be examples of galectin-3 influence on cancer cell metabolism. Of note there are many reports that point out metabolic reprogramming after SARS-CoV-2 infection.^101–104^

SARS-CoV-2 infection lead to many cardiovascular complications and postacute sequelae,^12,13,15,69,105,106^ including myocardial inflammation.^107–110^ Many reports point out the link between heart failure (HF) and cancer incidence.^106,111,112,112–118^ Systemic inflammatory signals are shown to be promoting cancer.^113^ There are also cases reported where cancer incidence occurred after COVID-19 infection. High level of STAT3 (signal transducer and activator of transcription-3) activation by IL-6 induces Epstein Barr virus (EBV) and Kaposi sarcoma-associated herpesvirus (KSHV) to enter lytic cycle.^119^ Case reports of an about 8-year-old boy diagnosed of having type B acute lymphocytic leukemia weeks after getting serologically tested positive for SARS-CoV-2 and Epstein-Barr virus^120^ and by Persaud et al.,^121^ are of interest. Acute B-cell lymphoblastic leukemia is also reported in two cases following SARS-CoV-2 infection.^122^ Bioinformatics and systems biology analyses also revealed common genes in gastric cancer and COVID-19.^123^ COVID-19 neutrophils were observed to be displaying neutrophil extracellular traps (NETs)^124^ and NETs were shown to be awakening dormant human breast cancer cells in mice^125^ suggesting a probable mechanism for COVID-19 influencing tumor relapse.^126^

Our study is based on bioinformatics and systems biology without experimental data. We took only three datasets for our analyses. More samples and experimental studies are necessary for the validation of our findings. We found hub genes and drugs targeting them that could be helpful in treating and (or) managing COVID-19 infections generally or in a tissue-specific manner. Eventhough there are some reports pointing towards cancer incidence post COVID-19, it is still unclear and our study also hints in this direction. In our analyses, increased expression of galactin3 was observed in the heart and the brain that has a positive influence in all the hallmarks of cancer. It could play a role in the incidence and progression of cancer after COVID-19 infection and this prompts in-depth studies in the incidence of cancer after SARS-CoV-2 infection. This could help in the inclusion of early cancer detection protocols in the health care system after SARS-CoV-2 infection for better prognosis in the unfortunate event of cancer incidence.

## Supporting information Data

Data 1: Supporting Information Data 1_All interacting proteins

Data 2: Supporting Information Data 2_Hub genes

Data 3: Supporting Information Data 3_Hub gene drug targets

Data 4: Supporting Information Data 4_Common and unique pathways

Data 5: Supporting Information Data 5_DEG biological pathways

Data 6: Supporting Information Data 6_Common and unique miRNA

Data 7: Supporting Information Data 7_Brain miRNET miR gene

Data 8: Supporting Information Data 8_Brain miRNA pathways

Data 9: Supporting Information Data 9_Lung miRNET miR gene

Data 10: Supporting Information Data 10_Lung miRNA pathways

Data 11: Supporting Information Data 11_miRNET heart genes

Data 12: Supporting Information Data 12_miRNAs_common_circulatory_brain_lung

Data 13: Supporting Information Data 13_Common miRNA_Circulatory_lung genes

Data 14: Supporting Information Data 14_Common miRNA_Circulatory_brain genes

Data 15: Supporting Information Data 15_Pathway list of 5 genes common in lung and brain

### CRediT authorship contribution statement

**Shrabonti Chatterjee:** Methodology, Writing – original draft, Validation, Formal analysis, Visualization. **Joydeep Mahata**: Methodology, Writing – original draft, Validation, Formal analysis, Visualization. **Suneel Kateriya:** Writing – review & editing, Supervision. **Gireesh Anirudhan:** Conceptualization, Methodology, Writing – review, editing and rewriting original draft, Visualization, Supervision.

DECLARATION OF COMPETING INTEREST The authors declare that they have no known competing financial interests or personal relationships that could have appeared to influence the work reported in this paper.

### Declaration of Competing interest

On behalf of all authors, the corresponding author states that there is no conflict of interest.

## Supporting information

Supporting Information Data 1_All interacting proteins

Supporting Information Data 2_Hub genes

Supporting Information Data 3_Hub gene drug targets

Supporting Information Data 4_Common and unique pathways

Supporting Information Data 5_DEG biological pathways

Supporting Information Data 6_Common and unique miRNA

Supporting Information Data 7_Brain miRNET miR gene

Supporting Information Data 8_Brain miRNA pathways

Supporting Information Data 9_Lung miRNET miR gene

Supporting Information Data 10_Lung miRNA pathways

Supporting Information Data 11_miRNET heart genes

Supporting Information Data 12_miRNAs_common_circulatory_brain_lung

Supporting Information Data 13_Common miRNA_Circulatory_lung genes

Supporting Information Data 14_Common miRNA_Circulatory_brain genes

Supporting Information Data 15_Pathway list of 5 genes common in lung and brain

## Acknowledgements

Ms. Shrabonti Chatterjee is a recipient of Non – National Eligibility Test (Non – NET) fellowship from University Grants Commission (UGC), Government of India. Mr. Joydeep Mahata is a recipient of DST – INSPIRE (Department of Science and Technology – Innovation in Science Pursuit for Inspired Research) Post – Graduate fellowship from Department of Science and Technology (DST), Government of India. We thank the above funding agencies for their financial support.

## Notes

### Competing Interest Statement

The authors have declared no competing interest.

### Summary of Updates

Discussion part is enhanced to show the significance of galectin-3 influence in cancer. One figure is modified to show gene regulation.

